# Astroglial CD38 regulates social memory and synapse formation through SPARCL1 in the medial prefrontal cortex

**DOI:** 10.1101/2021.12.23.474051

**Authors:** Tsuyoshi Hattori, Stanislav M Cherepanov, Ryo Sakaga, Jureepon Roboon, Dinh Thi Nguyen, Hiroshi Ishii, Mika Takarada-Iemata, Takumi Nishiuchi, Takayuki Kannon, Kazuyoshi Hosomichi, Atsushi Tajima, Yasuhiko Yamamoto, Hiroshi Okamoto, Akira Sugawara, Haruhiro Higashida, Osamu Hori

## Abstract

Social behavior is essential for the health, survival and reproduction of animals, yet the role of astrocytes in social behavior is largely unknown. CD38 is critical for social behaviors by regulating oxytocin release from hypothalamic neurons. On the other hand, CD38 is most abundantly expressed in astrocytes especially in the postnatal cortex, and is important for astroglial development. Here, we demonstrate that astroglial CD38 plays a pivotal role in the social behavior. Selective deletion of CD38 in postnatal astrocytes, but not in adult astrocytes, specifically impaired social memory without any other behavioral abnormalities. Morphological analysis revealed reductions in spine numbers, mature spines and excitatory synapse numbers in the pyramidal neurons of the medial prefrontal cortex (mPFC) due to deletion of astroglial CD38 in the postnatal brain. Astrocyte-conditioned medium (ACM) of CD38 KO astrocytes reduced synaptogenesis of cortical neurons by reducing extracellular SPARCL1, a synaptogenic protein. Finally, the release of SPARCL1 from astrocytes is regulated by CD38/cADPR/calcium signaling. Our data indicate that astroglial CD38 developmentally regulates social memory and neural circuit formation in the developing brain by promoting synaptogenesis through SPARCL1.

## Introduction

Social behavior means activities involving at least two individuals of the same species, which are critical for animals to survive and reproduce (1). Neural circuits which mediates various social behaviors such as aggression, mating, and parenting have been identified (1). Social memory plays a pivotal role in social behaviors, which is the ability to recognize a familiar or novel conspecific (2). It has been shown that neurons in the medial prefrontal cortex and the ventral CA1 and dorsal CA2 regions in the hippocampus are essential for social memory (3–6). On the other hand, poor social function is a prominent feature of several neuropsychiatric disorders, such as autism spectrum disorder (ASD) (7, 8). Genomic driven models of ASD display abnormal social behaviors together with abnormalities in formation, function and maintenance of neuronal synapses. *Fmr1* (fragile X mental retardation 1) knockout mice have shown to present abnormalities in social interaction, dendritic morphology and synaptic protein synthesis (9, 10). *Mecp2+/-* females, which are used to model Rett syndrome, exhibit reduced spine numbers and size of the cell bodies of neurons as well as abnormal social approach behaviors (11–14). A number of studies have indicated the importance of synaptic function in neural circuits that mediate social behaviors in various brain regions.

Astrocytes are the most abundant type of glial cells in the brain and physically interact with neurons, provide metabolic energy and neurotrophic factors, buffer extracellular ions, and modulate information processing and signal transmission (15). They influence synapse formation, maintenance and plasticity by releasing various molecules (16, 17), contributing to neural circuit formation, maturation and stabilization. Astrocytes affect cognitive processing including learning, memory and emotionality by regulating the synaptic connectivity of the neural networks (18, 19). These studies indicate that astrocytes play important roles in neuronal development and cognitive behaviors, however, the involvement of astrocytes in social behavior is largely unknown.

CD38 is a multifunctional transmembrane protein that possesses ADP-ribosyl cyclase activity, which produces cyclic ADP–ribose (cADPR) from nicotinamide adenine dinucleotide (NAD^+^), and releases Ca^2+^ from intracellular stores (20–22). CD38 has been shown to promote the secretion of oxytocin from hypothalamic neurons, affecting social behaviors (23). CD38 is dominantly expressed in astrocytes in the developing cortex, and astroglial CD38 promotes the differentiation of oligodendrocytes non-cell-autonomously (24). These studies suggest that astroglial CD38 plays a pivotal role in brain development through glia-glia crosstalk. However, the roles of astroglial CD38 in neural circuit formation and social behavior remains unknown.

In this study, we assessed the role of astroglial CD38 in social behavior and neuronal development. Selective deletion of astrocytic CD38 in the postnatal brain specifically impaired social memory and reduced excitatory synapse formation in the medial prefrontal cortex (mPFC). Using an *in vitro* culture system, CD38 KO astrocyte-conditioned medium (ACM) decreased synapse formation in cortical neurons, which was recovered by the addition of SPARCL1 protein, a synaptogenesis-promoting protein. Finally, secretion of SPARCL1 was regulated by CD38/cADPR/calcium signaling in astrocytes.

## Results

### Deletion of astroglial CD38 in postnatal brain impairs social memory

We previously showed that CD38 expression peaked from postnatal day 14 (P14) to P28 in the cortex and was most abundantly expressed in astrocytes during that period (24). To investigate the role of astroglial CD38 in social behavior, we generated conditional knockout mice in which *Cd38* gene was selectively deleted in the astrocytes of the postnatal brain using *CreER^T2^-LoxP* recombination technology. Because GLAST-CreER^T2^-mediated recombination was widely used to express transgenes and delete genes in astrocyte *in vivo* (25–27), homozygous *Cd38^flox/flox^* mice were crossed with *Glast^CreERT/+^* mice to generate *Cd38^flox/flox^;Glast^CreERT2/+^* mice. Tamoxifen was injected to *Cd38^flox/flox^;Glast^+/+^* or *Cd38^flox/flox^;Glast^CreERT2/+^* mice once a day for five consecutive days at P10–P14 (ctrl^P10^ or CD38 AS-cKO^P10^, respectively; Figure 1A). Western blot analysis demonstrated that CD38 expression in the cerebral cortex was substantially reduced in the brains of CD38 AS-cKO^P10^ mice compared to ctrl ^P10^ littermates at P21 (Supplementary Figure 1A and B).

**Figure 1.**
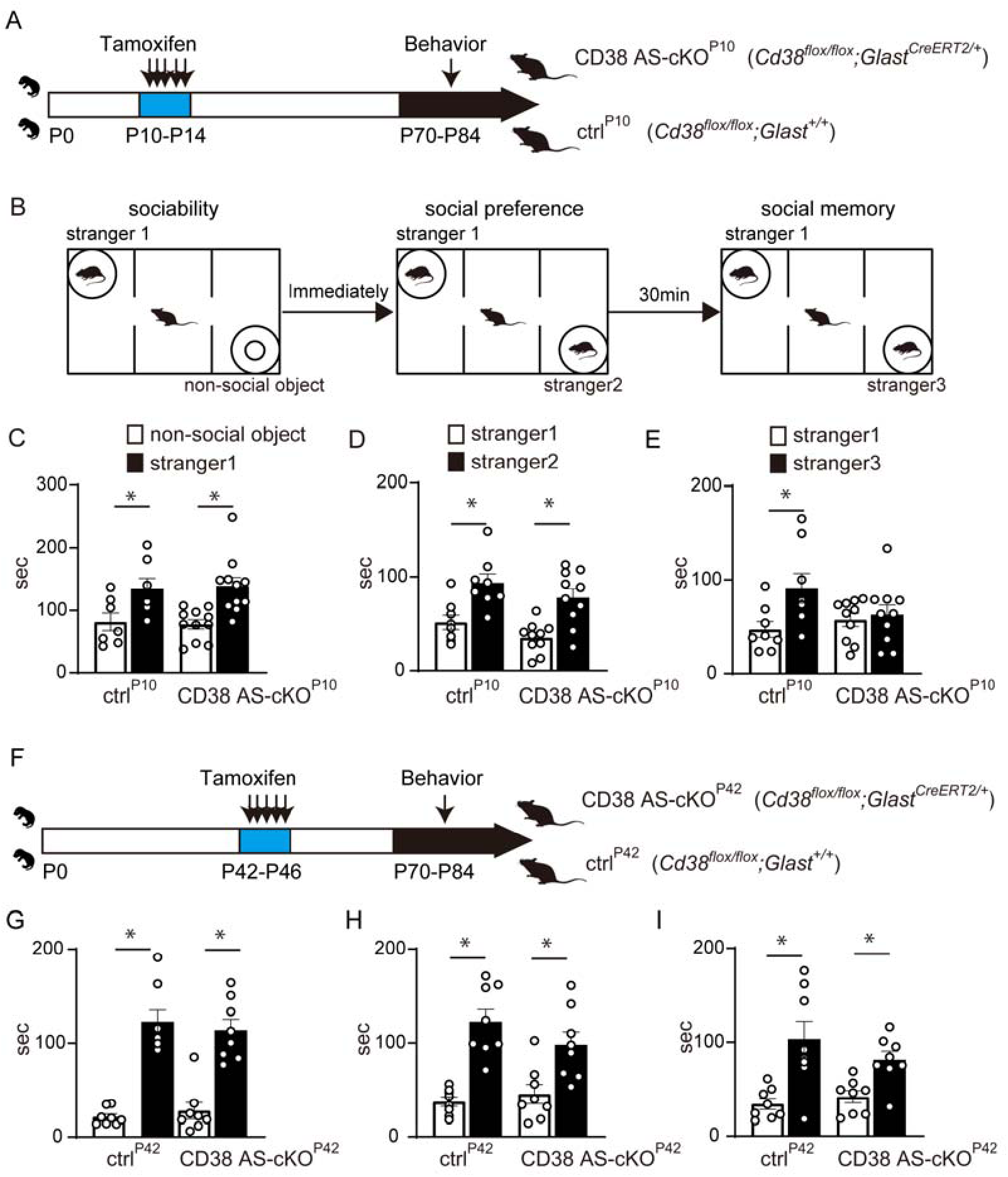
Deletion of astroglial CD38 in postnatal brain impairs social memory. (A, F) Paradigm for inducing recombination in which *Cd38^flox/flox^:Glast^CreERT2/+^* or *Cd38^flox/flox^:Glast^+/+^* mice receive 5 tamoxifen injections from P10 to P14 (ctrl ^P10^ or CD38 AS-cKO^P10^) or P42 to P46 (ctrl ^P42^ or CD38 AS-cKO^P42^). Behavior tests in these mice were performed from P70 to P84. (B) Schematic diagram of the three chamber test. (C, G) Sociability of mice in the three chamber test. All groups showed preference for the stranger mouse vs. an inanimate ball. (D, H) Social novelty in the three chamber test. All groups showed preference for the novel stranger vs. a familiar stranger. (E, I) Social memory in the three chamber test. CD38 AS-cKO mice^P42^ showed preference for a novel stranger vs a familiar stranger encountered 30 min ago (I), whereas CD38 AS-cKO^P10^ mice showed similar investigation time (E). *n* = 7 to 10 animals per genotype. Data represent means ± SEM. *P* values were determined by two-way ANOVA followed by Scheffe’s *F* test. \**p* < 0.05 between the non-social object and stranger 1, stranger 1 and stranger 2 or stranger 1 and stranger 3.

Next, we investigated astroglial development in this CD38 AS-cKO^P10^ mice. Consistent with our previous study using constitutive CD38 KO mice (24), the expression of astrocyte-specific molecules, GFAP, Cx43 and S100β was significantly decreased in the mPFC of CD38 AS-cKO^P10^ mice at P21 (Supplementary Figure 1A, C, D and K-N). Whereas the expression of NDRG2, another astrocyte-specific protein and number of NDRG2-positive astrocytes were not changed in these mice (Supplementary Figure 1A, E, O and P), indicating altered expression of several astrocyte-specific proteins without changing total number of astrocytes in CD38 AS-cKO^P10^ mice. On the other hand, unlike decreased myelination in constitutive CD38 KO mice (24), MAG and MBP expression, myelin proteins, was not altered in the cerebral cortex of CD38 AS-cKO^P10^ mice (Supplementary Figure 1A, H, Q and R). These results indicate that astrocyte-specific deletion of CD38 from P10 impairs postnatal development of astrocytes without affecting myelination.

Multiple studies have reported that constitutive CD38 KO mice exhibit impaired social behaviors (28–30). To investigate whether astrocytic CD38 is involved in social behavior, we conducted a three chamber social approach test of CD38 AS-cKO^P10^ mice, as shown in Figure 1B. In the first phase, the test mice were allowed to freely explore an object and stranger 1. Both ctrl ^P10^ and CD38 AS-cKO^P10^ mice showed a significant preference for exploring stranger 1 (Figure 1C). Immediately after the sociality test, the test mice were allowed to freely explore novel stranger 2 and the now-familiar stranger 1. Both ctrl ^P10^ and CD38 AS-cKO^P10^ mice demonstrated similar preference for social novelty (Figure 1D). After a 30 min interval, we performed the test again, with novel stranger 3 replaced from the stranger 2. Control mice spent more time investigating novel stranger 3 than the familiar stranger 1, whereas CD38 AS-cKO^P10^ mice showed similar investigation times between the familiar and stranger mouse (Figure 1E). These results indicate that CD38 AS-cKO^P10^ mice have deficit only in social memory. To investigate whether this deficit is attributed to abnormal brain development, we performed social behavior test of *Cd38^f/f^;Glast^+2/+^* or *Cd38^f/f^;Glast^CreERT2/+^* mice in which tamoxifen was injected from P42 to P46 (ctrl^P42^ or CD38 AS-cKO^P42^, respectively; Figure 1F). We confirmed significant reduction of CD38 protein expression in the mPFC of CD38 AS-cKO^P42^ (Supplementary Figure 1I and J). In these mice, all social behaviors, sociability, social preference, and social memory were similar to those of ctrl^P42^ mice (Figure 1G–I). Furthermore, constitutive CD38 KO mice have been reported to exhibit impaired parental, depression-like and hedonic behavior due to decreased plasma oxytocin levels (31). Thus, we performed a parental retrieval test, tail suspension test, sucrose preference test, elevated plus maze test, object recognition test and odor discrimination test. However, CD38 AS-cKO^P10^ mice did not show any significant differences in these behavioral tests nor in the plasma oxytocin levels compared to ctrl ^P10^ mice (Supplementary Figure 2A–J). These results indicate that deletion of astroglial CD38 specifically causes social memory deficits by disturbing postnatal brain development.

### Deletion of astroglial CD38 in postnatal brain impairs synapse formation in the medial prefrontal cortex

Previous studies have shown that the mPFC, ventral CA1 and dorsal CA2 regions in the hippocampus are essential for social memory (3, 5, 32). Furthermore, CD38 KO mice display prefrontal cortex dysfunctions such as impaired synaptic plasticity and excitation-inhibition balance (30). Thus, we investigated neuronal development in the mPFC of CD38 AS-cKO^P10^ mice. Golgi staining was performed to reveal changes in dendrites, spine density, and spine morphology in CD38 AS-cKO^P10^ mice at P70. The Number and total length of all dendrites and primary dendrites were quantified in the cortical pyramidal neurons of the mPFC. Consistent with a previous study on the motor and somatosensory cortex of constitutive CD38 KO mice (33), both indices of dendrites in CD38 AS-cKO^P10^ were not different from those of ctrl ^P10^ mice (Figure 2A–C). Next, spine densities and morphology were analyzed in the apical dendrites of cortical pyramidal neurons in the mPFC of CD38 AS-cKO^P10^ mice. Spine morphology was categorized into four groups, mushroom, branched, thin, and stubby spines. Mushroom and branched spines represent mature spines that have larger, more complex postsynaptic density, and are functionally stronger in response to glutamate. Thin spines and stubby spines represent immature spines that are flexible, rapidly enlarging or shrinking in brain development. Total spine densities were significantly reduced in apical dendrite of CD38 AS-cKO^P10^ mice (Figure 2D and E). Among the four spine types, mushroom and branched spines were significantly decreased in CD38 AS-cKO^P10^ mice. On the other hand, the number of thin and stubby spines did not change in CD38 AS-cKO^P10^ mice (Figure 2D and F). We further performed a morphological analysis of cortical neurons in the mPFC of constitutive CD38 KO mice. Constitutive CD38 KO mice as well as CD38 AS-cKO^P10^ mice also showed significantly decreased total spine and mature spine numbers (Supplementary Figure 3).

**Figure 2.**
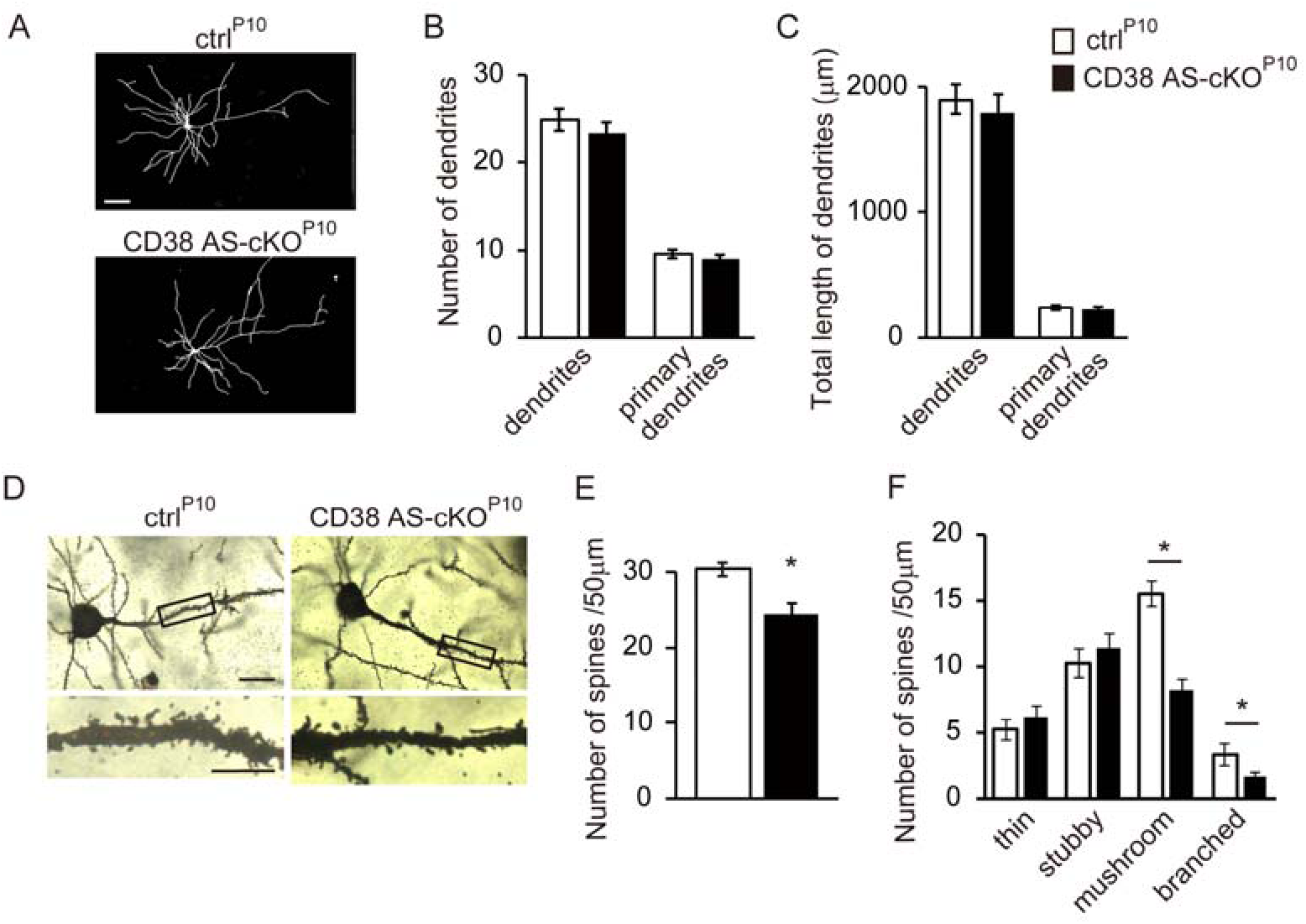
Spine morphology is altered in the mPFC pyramidal neurons in CD38 AS-cKO^P10^ mice. (A) Images of tracing of Golgi-stained pyramidal neurons in the medial prefrontal cortices of ctrl ^P10^ and CD38 AS-cKO^P10^ mice at P70. Scale bar = 20 μm. (B) Quantification of total number of dendrites and number of primary dendrites in ctrl ^P10^ and CD38 AS-cKO^P10^ mice. (C) Measurement of total length of dendrites and primary dendrites in ctrl ^P10^ and CD38 AS-cKO^P10^ mice. (D) Images of Golgi-stained pyramidal neurons in the mPFC of ctrl ^P10^ and CD38 AS-cKO^P10^ mice at P70. Scale bars = 20 μm (upper) and 5 μm (lower). (E) Quantification of the total number of spines on the apical dendrites of the ctrl ^P10^ and CD38 AS-cKO^P10^ mice. (F) The percentage of branched, mushroom, stubby, and thin spines in cortical apical dendritic spines. *n* = 6 cells (B and C) or 6 dendrites per animal, 4 animals per genotype. Data represent means ± SEM. *P* values were determined by Student’s unpaired *t*-test. \**p* < 0.05 vs. ctrl ^P10^.

We next assessed whether the number of synapses was decreased in the mPFC of CD38 AS-cKO^P10^ mice at P70 by immunostaining pre- and post-synaptic proteins, vesicular glutamate transporter 1 (VGlut1), and postsynaptic density protein 95 (PSD95), respectively. While there was no significant change in the density of VGlut1 immunoreactivity in CD38 AS-cKO^P10^ mice compared to littermate ctrl ^P10^ mice, there was a significant decrease in the density of PSD95 in CD38 AS-cKO^P10^ mice (Figure 3A–J). Consistent with the reduction in spine numbers and PSD95 expression, there was a significant decrease in the number of synapses defined as co-localized presynaptic VGlut1 and postsynaptic PSD95 immunoreactivity in CD38 AS-cKO^P10^ mice (Figure 3D, H andK). Further Western blotting analysis revealed decreased expression of PSD95 in CD38 AS-cKO^P10^ mice at P70 (Figure 3L–N). Constitutive CD38 KO mice also showed significantly decreased synapse numbers in the mPFC (Supplementary Figure 4), suggesting that CD38 regulates synapse formation non-cell-autonomously. These results indicate that the deletion of astroglial CD38 decreased excitatory synapses in the mPFC.

**Figure 3.**
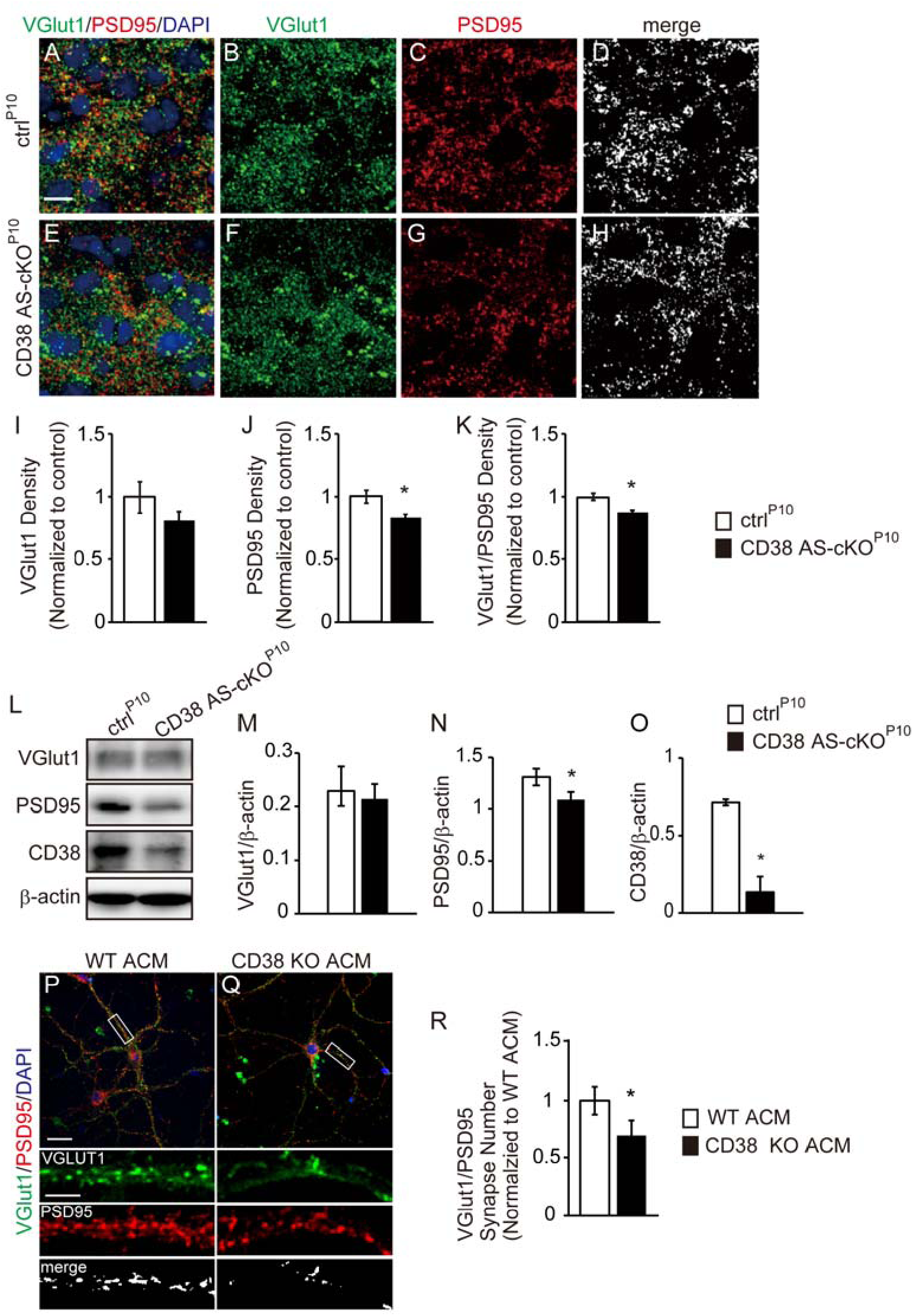
Astroglial CD38 promotes synapse formation of cortical neurons in the mPFC. (A–H) Immunohistochemistry for presynaptic marker VGlut1 (green) and postsynaptic marker PSD95 (red) in the mPFC of ctrl ^P10^ and CD38 AS-cKO^P10^ mice at P70. Images are single planes from confocal z-stacks. The VGlut1 and PSD95 channels are separated in panels B, C, F and G. Panels D and H are colocalized VGlut1 and PSD95 puncta. Nuclei were counterstained with DAPI. Scale bar = 10 μm. (I–K) Quantification reveals a significant decrease of PSD95 expression and synapses (colocalization of VGlut1 and PSD95) in the mPFC within CD38 AS-cKO^P10^ mice as compared to littermate ctrl ^P10^. *n* = 4 animals per genotype. (L) Western blot analyses of VGlut1, PSD95, CD38 and β-actin in the mPFC of ctrl ^P10^ and CD38 AS-cKO^P10^ mice at P70. (M) Bar graphs depict the relative optical density of VGlut1 and PSD95 normalized to the loading control β-actin. *n* =4 animals per genotype. Data represent means ± SEM. *P* values were determined by Student’s unpaired *t*-test. \**p* < 0.05 vs. ctrl ^P10^. (P, Q) Representative images of cortical neurons cultured for 14 days in vitro (DIV) in astrocyte conditioned medium (ACM) of WT or CD38 KO mice. Insets show individual channels for VGlut1 (green) and PSD95 (red) staining, as well as the merged image. Nuclei were counterstained with DAPI. Scale bar = 20 µm (main image) and 10 µm (inset). (R) Quantification reveals a significant decrease of synapses in cortical neurons cultured in CD38 KO ACM. *n* = 20 cells per condition from 3 independent culture. Data represent means ± SEM. *P* values were determined by Student’s unpaired *t*-test. \**p* < 0.05 vs. WT ACM.

### Astroglial CD38 promotes synapse formation of cortical neurons by secreted molecules

Astrocyte-conditioned medium (ACM) has a profound enhancement effect on synapse formation in neurons (17). To determine whether astroglial CD38 affects synaptic formation by secreted molecules, primary cortical neurons were cultured with WT- or CD38 KO-ACM from 4 to 14 DIV. Synaptic density was evaluated by immunostaining with anti-VGlut1 and -PSD95 antibodies. The number of synapses defined as colocalization of VGlut1 and PSD95 was significantly lower in neurons with CD38 KO-ACM than in those with WT-ACM (Figure 3P–R). On the other hand, there was no difference in the length of dendrites or viability between neurons cultured with WT- and CD38 KO-ACM (Supplementary Figure 5A–G). These results indicate that deletion of astroglial CD38 suppresses synapse formation through extracellular molecules.

### Astroglial CD38 promotes synapse formation through SPARCL1

To identify the extracellular molecules responsible for the decreased synapses in the mPFC of CD38 AS-cKO^P10^ mice and primary neurons treated with CD38 KO-ACM, we performed proteomic analysis of WT- and CD38 KO-ACM. A total of 118 proteins were identified by LC-MS/MS analyses of WT-ACM and CD38 KO-ACM. Proteins with an abundance ratio of (CD38 KO ACM) / (WT ACM) with 1.5 > or < 0.67 and abundance ratio variability (%) < 15 were subjected to gene ontology (GO) analysis. This analysis identified five extracellular proteins (Table 1). Among these proteins, SPARC-like protein 1 (SPARCL1) promotes synapse formation through its interactions with presynaptic NRX1a and postsynaptic NL1B (34). Proteomics analysis showed that SPARCL1 in CD38 KO-ACM was reduced to 58.3 ± 6.6 % of that in WT-ACM (Table 1). We further examined the amount of intracellular and extracellular SPARCL1 and another secreted protein, brain-derived neurotorophic factor (BDNF) in WT and CD38 KO astrocytes by Western blotting analysis of ACM and cell lysates of these cells. Although BDNF levels in ACM and cell lysate were not different between WT and CD38 KO astrocytes, both extracellular and intracellular SPARCL1 levels in CD38 KO astrocytes were significantly lower than those in WT astrocytes (Figure 4A-C). Next, we investigated SPARCL1 and CD38 expression in the mPFC of the developing brain. Consistent with a previous studies in the cortex (24, 35), expression of SPARCL1 and CD38 was developmentally regulated in the postnatal mPFC and peaked at P21 to P28 (Figure 4D-F), and SPARCL1 expression is largely restricted to S100β-positive astrocytes in the mPFC at P21 (Figure 4G and H). We found significantly decreased expression of SPARCL1 in astrocytes of CD38 AS-cKO^P10^ mice compared with that of ctrl^P10^ mice at P21 (Figure 4G and H). On the other hand, SPARCL1 expression in NeuN-positive neurons was not altered in CD38 AS-cKO^P10^ mice (Figure 4H). Since it has been reported that astroglial SPARCL1 promotes excitatory synapse formation in cultured neurons (36, 37), we first examined effect of SPARCL1 knockdown in cultured astrocytes on synaptogenesis. Transfection of SPARCL1 siRNA strongly reduced SPARCL1 protein both in cell lysate and ACM of astrocytes (Supplementary Figure 6A). ACM of SPARCL1 siRNA-transfected cells significantly decreased synapse formation of cortical neurons (Supplementary Figure 6B-D). Next, to confirm the involvement of SPARCL1 in CD38-mediated synaptogenesis, we evaluated synaptogenesis in neurons treated with recombinant SPARCL1 protein in CD38 KO-ACM. SPARCL1 partially recovered the decreased number of synapses in neurons cultured in CD38 KO-ACM (Figure 4I and J). These results demonstrate that CD38 induces excitatory synapse formation at least by promoting the release of SPARCL1 from astrocytes.

**Figure 4.**
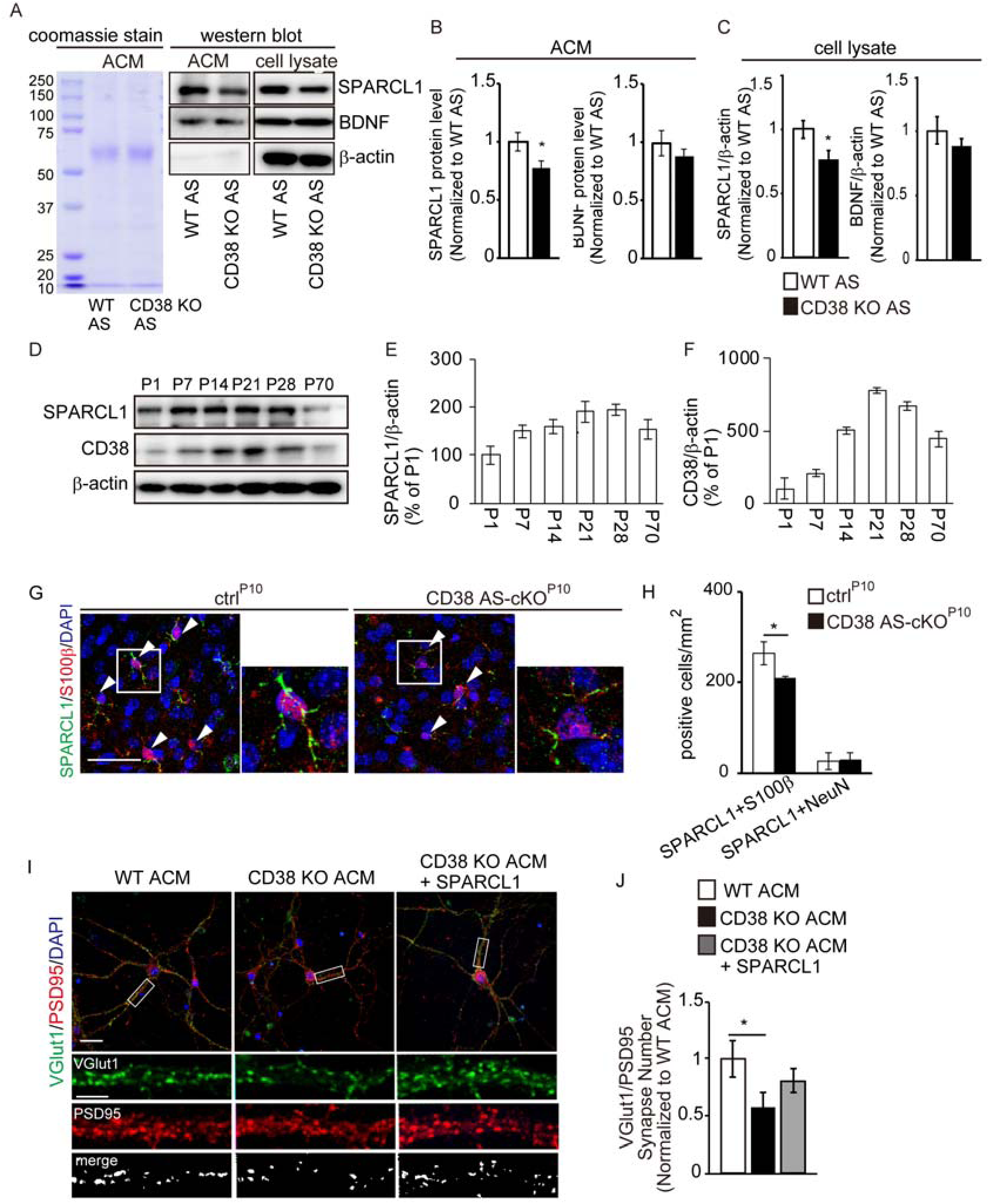
Astroglial CD38 promotes synapse formation through SPARCL1. (A) Coomassie stain shows the total protein composition of ACM of WT of CD38 KO astrocytes. Representative Western blot images of SPARCL1, BDNF and β-actin protein expression in ACM and cell lysates derived from WT or CD38 KO astrocytes. (B and C) SPARCL1 and BDNF level in ACM and cell lysates derived from WT or CD38 KO astrocytes. Protein levels in ACM and lysates were normalized to the total protein and the loading control β-actin, respectively. n = 6 independent culture per condition from 6 animals. Data represent means ± SEM. *P* values were determined by Student’s unpaired *t*-test. \**p* < 0.05 vs. WT AS. (D) Representative Western blot images of SPARCL1, CD38 and β-actin protein expression in the postnatal mPFC of WT mice. (E and F) Bar graphs depict the relative optical density of SPARCL1 and CD38 normalized to the loading control β-actin. *p☐<☐0.05 versus P70, n☐=☐3 animals. (G) Immunohistochemistry for SPARCL1 (green) and S100β (red) in the mPFC of ctrl ^P10^ and CD38 AS-cKO ^P10^ mice at P21. Arrowheads indicate SPARCL1/S100β-double positive cells. Nuclei were counterstained with DAPI. Scale bar =50 μm. (H) Bar graphs depict the number of SPARCL1/S100β- or SPARCL1/NeuN-double positive cells in the mPFC of ctrl ^P10^ and CD38 AS-cKO ^P10^ mice at P21. n = 4 animals per genotype. Data represent means ± SEM. *P* values were determined by Student’s unpaired *t*-test. \**p* < 0.05 vs ctrl ^P10^. (I) Representative images of cortical neurons cultured for 14 DIV in WT ACM, CD38 KO ACM and CD38 KO ACM with SPARCL1 proteins. Insets show individual channels for VGlut1 (green) and PSD95 (red) staining, as well as the merged image. Nuclei were counterstained with DAPI. Scale bar = 20 µm (main image) and 10 µm (inset). (J) Quantification reveals partially recovered synapses in cortical neurons cultured in CD38 KO ACM with SPARCL1 protein (80nM) compared with those cultured in CD38 KO ACM. *n* = 40 cells per condition from 3 independent culture. Data represent means ± SEM. *P* values were determined by one-way ANOVA followed by Tukey-Kramer test. \**p* < 0.05 vs. WT ACM.

**Table 1.** The proteomics of conditioned media from WT or CD38 KO astrocytes. The table shows extracellular proteins with an abundance ratio of (CD38 KO ACM) / (WT ACM), with 1.5 > or < 0.67, and an abundance ratio variability (%) < 15. n = 2 independent cultures from 2 animals per genotype.

| Uni ac No. | protein | KO ACM/WT<br>ACM (%) | Variation (%) |
| --- | --- | --- | --- |
| Q6GU68 | Immunoglobulin superfamily containing leucine-rich repeat protein | 38.9 | 10.02 |
| Q91WP6 | Serine protease inhibitor A3N | 43.6 | 6.31 |
| P50608 | Fibromodulin | 53.1 | 0 |
| **P70663** | **SPARC-like protein 1** | **58.3** | **6.61** |
| Q61702 | Inter-alpha-trypsin inhibitor heavy chain H1 | 59.4 | 12.94 |

CD38 catalyzes the synthesis of a calcium messenger cADPR, which releases Ca^2+^ from intracellular stores (38), and CD38/cADPR/Calcium signaling is involved in astrocytic production of extracellular mitochondria (39). Thus, we assessed the role of this pathway in the release of extracellular SPARCL1 from astrocytes. We used 8-Bromo-cADPR, a cADPR antagonist, 2APB, an IP_3_ inhibitor, and dantrolen, a ryanodine receptor inhibitor, to block cADPR/calcium signaling (Figure 5A). All these compounds significantly decreased the release of SPARCL1 from astrocytes (Figure 5B and C). On the other hand, they did not change the expression of intracellular SPARCL1 in astrocytes (Figure 5B and D), indicating that CD38/cADPR/calcium signaling is involved in the release of SPARCL1 from astrocytes. Finally, we performed bulk RNA-sequencing (RNA-seq) on cultured astrocytes from WT and CD38 KO mice (n = 2 independent cultures per genotype.). Our results showed that deletion of CD38 changed the gene expression profile in astrocytes (Figure 6A). We performed Kyoto Encyclopedia and Genes and Genomes (KEGG) pathway analysis on 262 decreased and 397 increased genes in CD38 KO astrocytes (log_2_ fold change < -0.57 or >0.57; FPKM>3; FDR < 0.05) from the RNA-seq data, ranked by *p* value (Figure 6B). We found calcium signaling pathway was significantly down-regulated in CD38 KO astrocytes (Figure. 5C). On the other hand, most of the pathways up-regulated in CD38 KO astrocytes were related with inflammatory response (Figure 6D).

**Figure 5.**
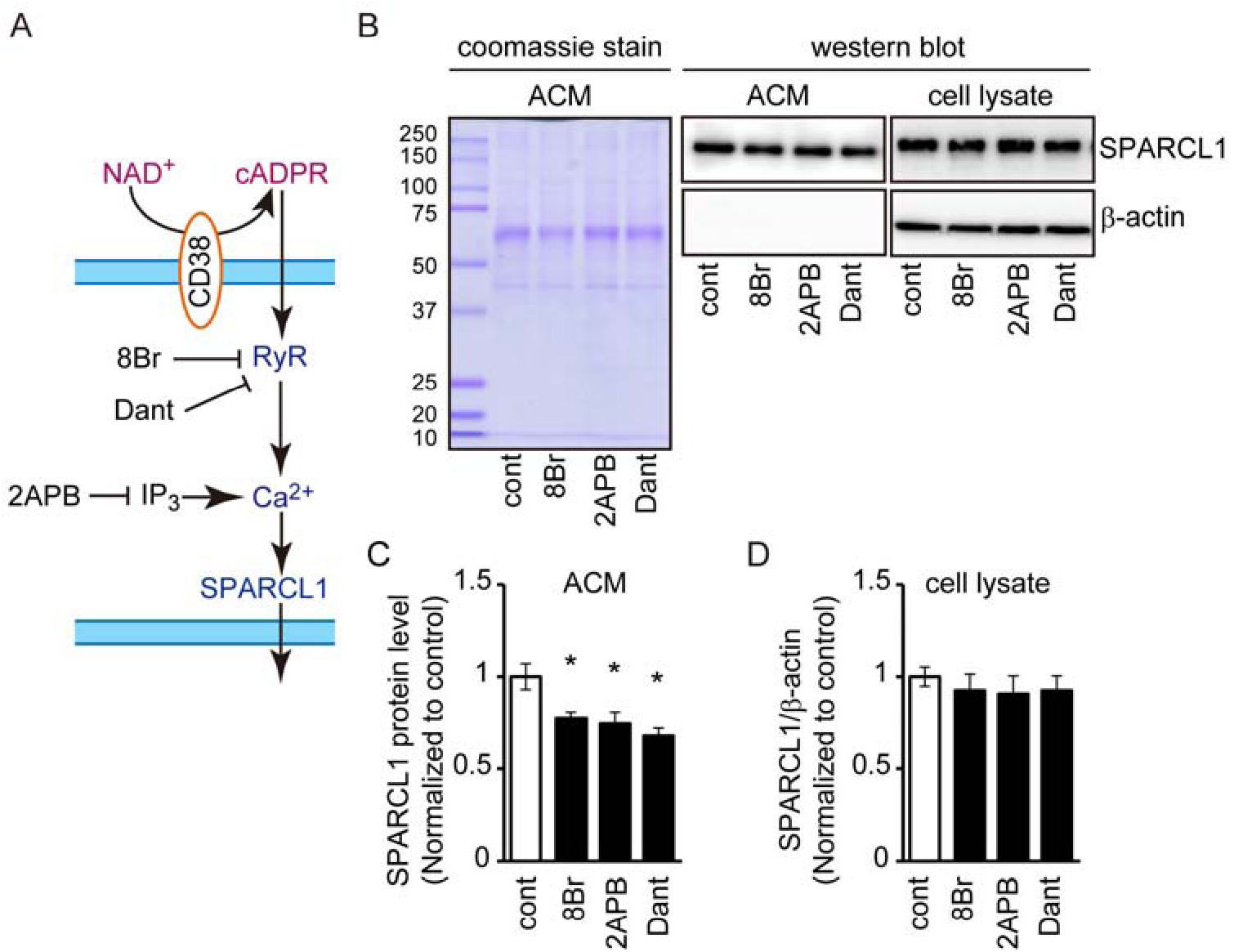
cADPR/calcium signaling regulates release of SPARCL1 from astrocytes. (A) cADPR produced by the catalytic action of CD38 from NAD^+^ acts as a calcium releasing second messenger via ryanodine receptors (RyR). 8-Bromo-cADPR (8Br): a cADPR antagonist, 2APB: a IP_3_ inhibitor and dantrolen (Dant): a ryanodine receptor inhibitor that blocks cADPR/calcium signaling (B) Coomassie stain shows the total protein composition of ACM. Representative Western blot images of SPARCL1 and β-actin protein expression in ACM and cell lysates from WT astrocytes treated with 8-Br-cADPR (50μM) or Dantrolen (20μM). (C and D) SPARCL1 level in ACM and cell lysate derived from WT or CD38 KO astrocytes. Protein levels in ACM and lysates were normalized to the total protein and the loading control β-actin, respectively. n = 5 independent culture per condition from 5 animals. Data represent means ± SEM. *P* values were determined by one-way ANOVA followed by Tukey-Kramer test. \**p* < 0.05 vs control.

**Figure 6.**
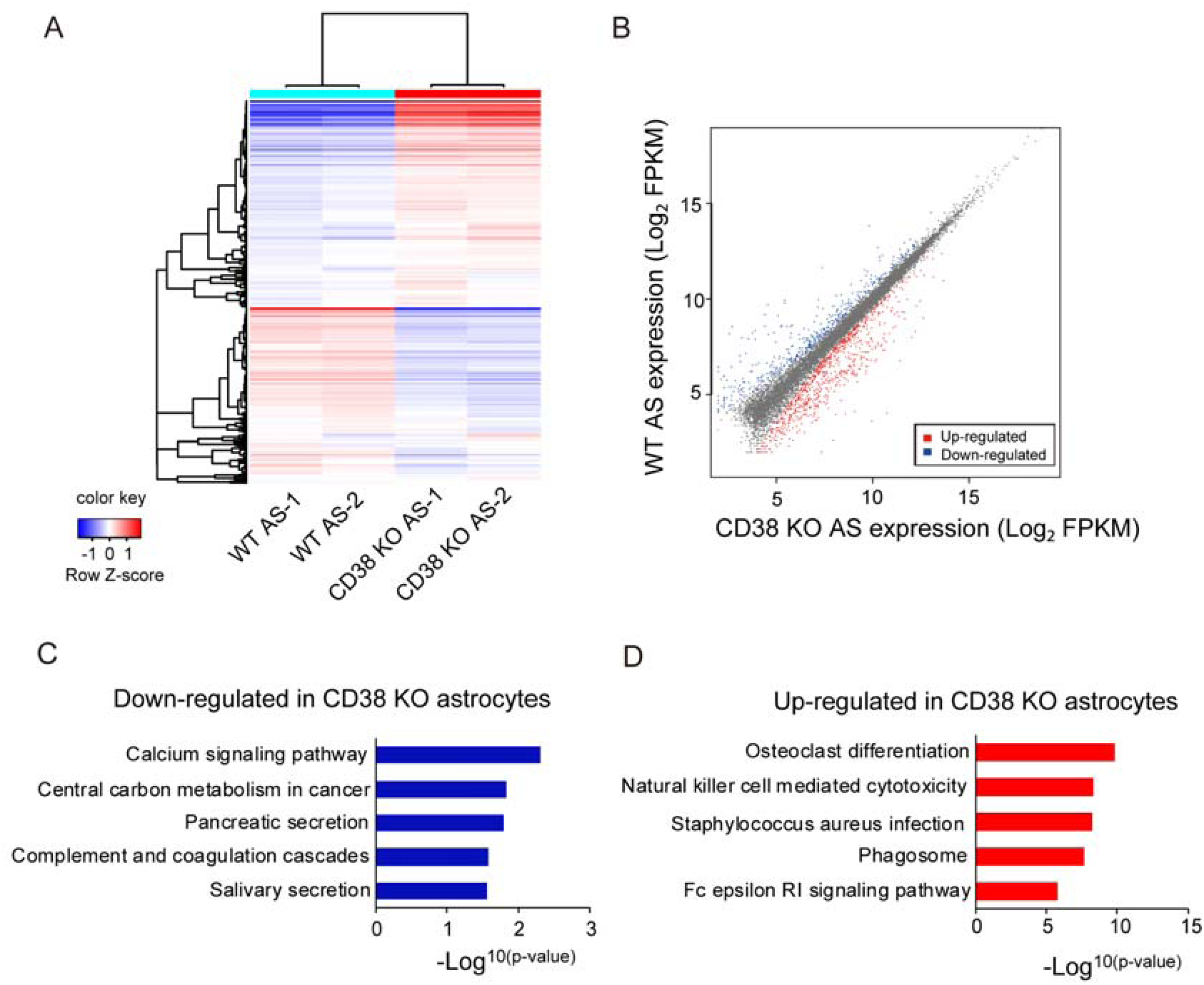
Transcriptome analyses reveals altered calcium signaling pathway in CD38 KO astrocytes. (A) Heat maps of RNA-seq data from WT and CD38 KO cultured astrocytes. Genes and samples were clustered using correlation distance with complete linkage. (B) Scatter plot of transcripts up-regulated (red dots) and down-regulated (blue dots) in WT and CD38 KO astrocytes (FDR<0.05, Fold change>1.5), n = 2 independent cultures from 2 animals per genotype. (C) Graph depicting top ranked KEGG terms by gene ontology analysis of down- and up-regulated genes in CD38 KO astrocytes. Significance of enrichment in KEGG pathway analyses was calculated using the p value function given from a modified Fisher’s exact test by the DAVID database.

## Discussion

In the current study, we clarified the role of astroglial CD38 in social behavior. Unexpectedly, we found that astroglial CD38 regulated specifically social memory in the postnatal brain and synapse formation in the mPFC. CD38 regulates SPARCL1 expression and secretion in astrocytes, which contributes to synaptogenesis in cortical neurons. Finally, SPARCL1 secretion from astrocytes is regulated by CD38/cADPR/calcium signaling. These results demonstrate that astroglial CD38 plays a pivotal role in social memory by regulating neural circuit formation in the postnatal brain.

### Astrocytes and social memory

Multiple studies have shown the involvement of astrocytes in cognitive processes such as learning, memory, and attention by manipulating astrocyte-specific molecules (16). For the first time, we demonstrated the involvement of astroglial CD38 in social memory. Although deletion of astroglial CD38 from P10 completely impaired social memory, its deletion from P42 did not cause social memory deficits (Figure 1). These results indicate that astroglial CD38 plays a role in neural circuit formation in social memory by promoting synapse formation. Synapse formation in the cortex starts from the first to third week of postnatal cortical development, which is concurrent with astrocyte development processes including division, expansion, and process outgrowth (40). Notably, synaptic numbers and total spine numbers were decreased and spine maturation was impaired in the mPFC of CD38 AS-cKO^P10^ mice (Figure 2 and 3). Impaired synapse formation was observed in the excitatory pyramidal neurons of the mPFC (Figure 3). Consistent with our findings of the behavioral phenotypes, the pyramidal neurons in the mPFC project axons to the nucleus accumbens, which does not affect social investigation but is critical for social memory (6).

Therefore, deletion of astroglial CD38 may impair synapse formation in neurons from the mPFC to the Nucleus Accumbens. However, astroglial CD38 is not limitedly expressed in the mPFC and is expressed in other brain areas such as the cortex, cerebellum, and hippocampus (24, 41). Thus, deletion of astroglial CD38 may affect synapse formation of neurons in the areas that are critical for social memory, such as the ventral CA1 and CA2 areas in the hippocampus (3, 5).

Previous studies have shown behavioral deficits in constitutive CD38 KO mice, including anxiety-like behaviors, locomotor activity deficits, and altered parental and social behaviors (23, 28, 31). However, CD38 AS-cKO^P10^ mice did not show any of these behavioral deficits and demonstrated social memory. In constitutive CD38 KO mice, plasma oxytocin levels were decreased and most of the behavioral deficits were recovered by oxytocin administration. These deficits are attributed to lower oxytocin levels by deletion of CD38 in oxytocin neurons in the hypothalamus. We found that plasma oxytocin levels did not change in CD38 AS-cKO^P10^ mice (Supplementary Figure 2J). Therefore, astroglial CD38 does not seem to affect oxytocin release from oxytocin neurons. A previous study has shown that constitutive CD38 KO mice exhibit learning and memory impairments in the Morris water maze, contextual fear conditioning test, and novel object recognition test (29).

However, the involvement of oxytocin in memory dysfunction has not yet been studied. Moreover, many studies have shown that astrocyte-specific molecules, including extracellular molecules such as IL-6, are involved in memory function (16, 42). Thus, astroglial CD38 may also be involved in learning and memory functions. Further studies are necessary to clarify the common and specific mechanisms of the regulation of social memory and other memory functions in neurons and astrocytes.

### Regulation of synapse formation by astroglial CD38

We identified SPARCL1 as a potential molecule for synapse formation in cortical neurons, which is regulated by astroglial CD38. This finding is congruent with previous studies demonstrating a role of SPARCL1 for promoting excitatory synapse formation (36, 37). Furthermore, we found astrocyte-specific expression of SPARCL1 in the postnatal mPFC (Figure 4H and I) as in the visual cortex (35). Therefore, SPARCL1 is thought to be involved in the CD38-mediated neural circuit formation of social memory at least in the mPFC. However, the addition of recombinant SPARCL1 did not fully recover the impaired synapse formation by CD38 KO ACM (Figure 4I and J). This suggests that other known or unknown synapse-affecting molecules are regulated by astroglial CD38. In addition to SPARCL1, we identified four other extracellular proteins that were downregulated in CD38 KO ACM (Figure 4A). Although these molecules, except the immunoglobulin superfamily containing leucine-rich repeat (ISLR), are reported to be expressed in astrocytes (43–45), the functions of these molecules that are related to neuronal development, including synapse formation, have not been reported. However, there is still a possibility of the involvement of these molecules in synapse formation. As with SPARCL1 (46), fibromodulin regulates extracellular matrix assembly, which contributes to glioma cell migration (44).

We found that blocking cADPR/calcium signaling decreased the secretion of SPARCL1 from astrocytes (Figure 5B and C). Furthermore, suppression of CD38 expression downregulates astrocytic cADPR/calcium signaling (38). Thus, the decreased extracellular SPARCL1 in CD38 KO ACM is likely attributed to the downregulation of this signaling pathway. Similarly, the CD38/cADPR/calcium pathway in astrocytes also regulates mitochondrial transfer to neurons in brain ischemia (39), which may be important for crosstalk between astrocytes and neurons by regulating the release of astrocyte-derived factors. Furthermore, intracellular SPARCL1 expression was significantly decreased in CD38 KO astrocytes (Figure 4B and D). In addition to the decreased release, this also contributed to the decreased extracellular SPARCL1 in CD38 KO astrocytes. However, blocking of the cADPR/calcium pathway did not decrease its expression (Figure 5D). In the present study, astrocytes were treated with a cADPR antagonist or calcium inhibitors for 48 h. Continuous long-term suppression of cADPR/calcium signaling may be necessary for downregulating SPARCL1 expression. Alternatively, elevation of NAD^+^ levels by deletion of CD38 may suppress SPARCL1 expression.

In conclusion, astroglial CD38 developmentally regulates social memory and neural circuit formation in the mPFC by promoting synaptogenesis through SPARCL1. The identification of a novel molecule that mediates social memory without affecting social preference may provide novel targets for the treatment of diseases with social memory deficits, such as autism.

## Materials and Methods

### Animals

All animal experiments were approved by the Animal Care and Use Committee of Kanazawa University (AP-194042). *Cd38^flox/flox^* mice were obtained from Akira Sugawara, Tohoku University. *Glast^CreERT2/+^* mice were obtained from Magdalena Götz, GSF-National Research Center for Environment and Health (47). For Cre-lox experiments, *Cd38^flox/flox^:Glast^CreERT2/+^* were crossed with *Cd38^flox/flox^:Glast^+/+^*. Male *Cd38^flox/flox^:Glast^CreERT2/+^* and *Cd38^flox/flox^:Glast ^+/+^*were injected with tamoxifen (20mg/kg; T-5648, Sigma-Aldrich, St. Louis, MO, USA) intraperitoneally once a day for 5 consecutive days from P10 or P42, a protocol previously demonstrated to induce efficient recombination (48). All mice were maintained by breeding to ICR mice. CD38 KO mice were generated as described previously (49).

### Three chamber test

The three chamber social apparatus was a (90 cm × 50 cm × 30 cm) plexiglass box, which was divided into three equal size compartments by two transparent partitions. At the floor level of each partition, there was a square opening (5 cm × 5 cm) located in the center, allowing access into each chamber. Two small, round wire cages were put in the diagonal corner of the apparatus for enclosing a stranger mouse or a similar size ball and two weighted bottles were placed on the top of the cages to prevent the test mice from climbing over the wire cages. The test was designed in accordance with previous study (50). The day before the test, all the test mice were habituated to the apparatus for 20 min with the two empty cages inside and all the stranger mice were habituated inside the wire cages for 20 min at a separate time. On the test day, after a 10-min habituation period, a stranger mouse was placed into one of the two wire cages while an inanimate ball was placed into the other. Then, the subject mouse was placed in the center, and allowed to freely explore the chamber for 5 min. The subject had the choice between a stranger mouse and an inanimate ball in this phase (sociability phase). Immediately after the sociability phase, the inanimate ball was replaced by a second stranger mouse, and the subject was allowed to freely explore the chamber for another 5 min. Thus, the subject would now have the choice between the mouse that the subject had already encountered and a new stranger mouse (preference for social novelty phase). After an interval of 30 min, the second stranger mouse was replaced by a third stranger mouse, and the subject was allowed to freely explore the chamber for 5 min again. Now it had the choice between familiar stranger 1 that it had already encountered 30 min before and the unfamiliar stranger 3 (social memory phase). Interaction time was scored when the subject was sniffing towards the cages within 2 cm. The location of the stranger 1 was alternated between tests. All the stranger mice used in the test were age- and gender-matched ICR mice. Researchers were blind to genotypes during experiments. Behavior was recorded on video-source and was analyzed using ANY-Maze behavioral tracking software (Stoelting Co., Wood Dale, IL, USA).

### Primary astrocyte culture

Astrocytes were isolated from the cerebral cortex of P1 to P3 neonatal mice following methods previously described (51). Briefly, cerebral cortices were harvested from neonatal mice. The brain tissue was then digested at 37°C in 4-(2-hydroxyethyl)-1-piperazineethanesulfonic acid (HEPES)-buffered saline containing Dispase II (383-02281, 2 mg/mL; Wako). Cells were plated in PDL-coated flasks (356537, Corning, Corning, NY, USA) in D-MEM (044-29765, WAKO) supplemented with 10% fetal bovine serum (172012, Sigma-Aldrich). After 10 days of cultivation, microglial cells and oligodendrocytes were removed by aspiration following 20 h shaking and the adherent cell population was collected after being detached using 0.05% trypsin-EDTA (15400054, Thermo Fisher Scientific). Collected cells were plated at 1 × 10^5^ cells/cm^2^ density. The next day the medium was changed in order to remove cell debris and loosely attached cells. Medium was changed every day 3days..

### Primary neuron culture

Primary cortical neurons were prepared from embryonic day 18 BL6 mice with nerve cell culture system 291-78001 (FUJIFILM Wako Chemicals Co. Ltd, Tokyo, Japan). 4 well chamber slides (177437, Thermo Fischer Scientific) were coated by incubating for 12 h with filter sterilized poly-L-lysine (P8920, Sigma-Aldrich; 100 μ g/ml) in borate buffer (boic acid 3.1g/l, borax 4.8g/l in water). Then, they were washed with sterile water and dried before use. Neurons were plated on the chamber slides at a density of 1 × 10^5^ cells per well and cultured in Neurobasal medium (21103049, Thermo Fischer Sicentific) with 5% FBS, 2% B27 supplement (A3582801, Thermo Fischer Scientific), 100ug/ml Penicillin-Streptomycin (168-23191, WAKO) and 0.5mM L-Glutamine (076-00521, WAKO).

### Primary neuron culture with conditioned medium from astrocyte cultures

Astrocytes were isolated from wild-type (WT) or CD38 KO neonatal mice as described above. After 9 days of cultivation, cells were replated in 10 cm diameter dishes at the density of 5 × 10^5^ cells per dish and cultured for 48 h. Then all of the media was replaced with Neurobasal medium supplemented with 2% B27 supplement and 0.5mM L-glutamine after washing with Phosphate-buffer saline (PBS) 3 times. The media were harvested 24 h after replacement, and filtrated by Millex-GP filter (SLGPR33RS, Merck Millipore) and centrifuged at 3000 g for 3 min to remove cellular debris. Primary neuron cultures were isolated as described above and plated on 4 well chamber slide coated with poly-L-lysine. Three days after plating, cells were cultured with 5 μ AraC (C6645, Sigma-Aldrich) for 24 h. Four days after plating, half of the media were replaced with ACM. At day nine, half of the medium was replaced with ACM again. The neurons were cultured with or without SPARCL1 recombinant protein (80 nM, ATGP3891, NKMAX) from day 8 to day 14 as previously done (34). Cells were fixed by 4% paraformaldehyde for immunostaining at day14.

### Golgi staining

Male mice (4 ctrl ^P10^, 4 CD38 AS-cKO^P10^, 4 WT and 4 CD38 KO) aged 10 weeks were used. Brains were removed and Golgi-Cox staining was performed using an FD Rapid GolgiStain Kit (PK401, FD NeuroTechnologies, Ellicott City, MD, USA) according to the manufacturer’s instructions. Unfixed brain samples were processed as previously reported (52). The coronal brain blocks were immersed in a solution of equal parts Solution A and B at room temperature for 2 weeks and then soaked in Solution C at 4°C for 48 h. After freezing with dry-ice powder, the brain samples were sliced into 250 µm pieces at -22°C using a cryostat microtome. Each frozen section was mounted with Solution C on a 0.5% gelatin-coated glass slide. Slides were allowed to dry naturally at room temperature and were then stained with a mixture (Solution D: Solution E: DW; 1:1:2) for 5 min, after which the slides were rinsed in distilled water twice for 4 min each. Slides were dehydrated in an ascending ethanol series and then sealed with Entellan (Merck, Darmstadt, Germany) through xylene. The preparations were observed in detail with a BZ-9000 digital microscope (Keyence Corporation, Osaka, Japan).

To assess dendritic morphology, low magnification (20× lens) images (Z-stack with 1 μm intervals) of pyramidal neurons, with cell bodies located in Layer II/III of the mPFC, were randomly captured. Captured images from ctrl ^P10^, CD38 AS-cKO ^P10^, WT and CD38 KO neurons were manually traced using Simple Neurite Tracer plug-in (SNT) from FIJI (53). Total length and number of all the dendrites and primary dendrites were measured with SNT.

To assess dendritic spine morphology, high magnification (63× lens) images (Z-stack with 0.2 μm intervals) of pyramidal neurons were captured. The density of spines was quantified by counting the number of spines on the following segments: dendritic segments (50-100 μm) of apical dendrites located farther than 50 μm from cell soma of ctrl ^P10^, CD38 AS-cKO ^P10^, WT and CD38 KO neurons. The spines were classified based on their morphology such as a mushroom type (diameter of the spine head was larger than that of the spine neck), thin type (diameter of the spine head was equal to that of the spine neck), stubby spines (no neck), and ramified spines (two heads).

### Western blot analyses

Conditioned media from astrocytes were collected at 5 DIV and filtrated with Millex-GP filter (SLGPR33RS, Merck). ACM was concentrated with Amicon Ultra 10kDa (UFC501008) following the manufacture’s protocol. Cells and brain samples (mPFC) were homogenized and solubilized in a buffer containing 1% NP40, 0.1% sodium dodecyl sulfate, and 0.2% deoxycholate. Lysates and ACM were boiled for 5 min with sodium dodecyl sulfate (SDS) sample buffer, subjected to SDS-polyacrylamide gel electrophoresis (SDS-PAGE) and proteins were transferred to polyvinylidene fluoride (PVDF) membranes. After blocking in 5% skimmed milk, the membrane was incubated for 12 h at 4°C with the primary antibody: anti-VGlut1(AB5905, Millipore, Burlington, 1:2000), anti-PSD95 (51-6900, Thermo Fischer Scientific, 1:500), anti-CD38 (AF4947, R&D systems, 1:300), anti-β-actin (013-24553, WAKO, 1:2000), anti-SPARCL1 (AF2836, R&D systems, 1:2000); anti-BDNF (ab108319, Abcam, 1:2000), anti-GFAP (G9269, SIGMA, 1:2000), anti-Cx43 (MAB3068, Merck Millipore, 1:200), anti-NDRG2 (sc-19468, Santa Cruz Biotechnology, 1:500) and anti-MAG (sc-15324, Santa Cruz Biotechnology, 1:500) . The membrane was incubated for 1 h at room temperature with anti-rabbit, mouse or goat horseradish peroxidase (HRP)-linked IgG (sc-2357, sc-516102, sc-2354, Santa Cruz Biotechnology, 1:5000).

Immunoreactivity was detected using an enhanced chemiluminescence system (ELLUF0100, Merck). Densitometric quantification was performed using ImageJ 1.52 software (https://imagej.nih.gov/ij/).

### Immunohistochemistry and synaptic puncta analysis

The brains were removed from the mice after perfusion with 4% paraformaldehyde (PFA). The brains were fixed with 4% PFA in PBS at 4°C overnight and then cryoprotected with 30% sucrose in PBS overnight. Cortical sections (20 µm-thick sections) were cut on a cryostat (Leica CM1950, Leica, Germany). Sections were washed and permeabilized in PBS with 0.3% Triton-X 100 three times at room temperature. Sections were blocked in 3% Bovine Serum Albumin (BSA, 015-21274, WAKO) in PBS for 1 hr at room temperature.

Primary antibodies against GFAP (1:2000), NDRG2 (1:500), VGlut1 (1:2000), PSD95 (1:500), S100β (S2532, Sigma-Aldrich, 1:300), NenN (MAB377, Millipore, 1:500), MBP (MAB386, Millipore, 1:300) and SPARCL1 (1:500) were diluted in 3% BSA containing PBS. Sections were incubated with primary antibodies overnight at 4°C. Alexa488- or Cy3-conjugated secondary antibodies (1:300) were used to visualize immunolabeling. Slides were mounted in Vectashield with DAPI and images were acquired on a laser scanning confocal microscope (Dragonfly; Andor, Beifast, Northern Ireland).

4 animals/genotype were stained with pre- (VGlut1) and post-synaptic (PSD95) marker pairs as described previously (35). Three independent coronal sections per each mouse, which contain the mPFC (Bregma -2.0--2.8 mm) were used for analyses. 5 μm thick confocal z-stacks (optical section depth 0.33 μm, 15 sections/z-stack, scanned area 0.04 mm^2^) of the layer II/III in the prelimbic region of the mPFC were imaged with a 63×/1.20 W HC PL APO CORR CS2 objective on a Dragonfly confocal laser scanning microscope.

Maximum projections of three consecutive optical slices (corresponding to 1 μm total depth) were generated from the original z-stack. A total of 5 confocal z-stacks per animal were analyzed for blind analysis using ImageJ 1.52 software. 1 μm thick maximum projections were separated into green and red channels and thresholded to detect discrete puncta without introducing noise. Density of thresholded pre and postsynaptic markers was measured using the Measure Particles function where a puncta size was defined for each marker (Vglut1 = 0.1-infinity; PSD95 = 0.1-infinity). The colocalization of puncta was quantified using the Image Calculator function applied to thresholded pre and postsynaptic images. The synapse and terminal densities were calculated by taking the total puncta area and dividing it by the total area of the field of view. Densities for ctrl^P10^ were averaged, then all image values were converted to ratio to the calculated control^P10^ average.

### Immunocytochemistry

Immunofluorescent staining of primary cultures was performed as previously described (54). Briefly, cells were fixed with 4% PFA for 10 min, permeabilized with 0.1% Triton-X for 10 min, and blocked with 3% BSA for 1 h at room temperature. After overnight incubations with antibodies against VGlut1 (1:3000) and PSD95 (1:1000) at 4°C, the cells were incubated with Alexa-488 and Cy3-conjugated secondary antibodies and DAPI for 1 h at room temperature. Imaging was performed at 63× magnification on a laser scanning confocal microscope (Dragonfly). Synaptic puncta were counted along well-isolated primary dendrites (100 μm dendritic segments) using a 63× lens on a fluorescence microscope. The number of co-localized puncta was calculated by ImageJ 1.52 as described above. All image values were converted to ratio to the calculated WT ACM average.

### Comparative Shotgun Proteomics using nano-liquid chromatography mass spectrometry

After the WT and CD38 KO astrocytes reached confluency at 7 DIV, cells were washed three times with PBS and 10 ml of neurobasal medium was added. After 24 h, astrocyte conditioned medium and control neurobasal medium were collected and filtrated with Millex-GP filter. The protein pellets were resuspended in the mixed buffer of 6 M Urea and 50 mM triethylammonium bicarbonate (TEAB) in pH 8.5. After quantification of protein concentration by BCA method, all proteins (10 µg) were adjusted to a final volume of 10 µL. Comparative shotgun proteomics were performed as previously described (55).

Briefly, these proteins were reduced, by tris(2-carboxyethyl)phosphine (TCEP), alkylated by iodoacetamide (IAA), and digested by trypsin. The trypsin-digested peptides were purified and separated on the LC system (EASY-nLC 1200; Thermo Fisher Scientific). The peptide ions were detected using MS (Orbitrap QE plus MS; Thermo Fisher Scientific). The MS/MS searches were carried out using SEQUEST HT search algorithms against the Mus Musculus (Swiss prot. Tax ID 10090) protein database (2017-6-7) using Proteome Discoverer (PD) 2.2 (Version 2.2.0.388; Thermo Fisher Scientific). Label-free quantification was also performed with PD 2.2 using precursor ions quantifier nodes. Normalization of the abundances was performed using total peptide amount mode. Among identified proteins in WT and CD38 KO ACM, Proteins with abundance Ratio: (CD38 KO ACM) / (WT ACM) with 1.5 > or < 0.67 and abundance Ratio Variability (%) < 15 were subjected to following a gene ontology (GO) analysis using the online version of DAVID Bioinfomatics Resources 6.8 (https://david.ncifcrf.gov/home.jsp). A functional annotation analysis was performed with the gene ontology tool (GOTERM_CC_ALL) using the UniProt Accesion numbers of the proteins and extracellular proteins were shown in Table 1. The list of the identified proteins in WT and CD38 KO ACM were shown in Supplementary Material 2.

### RNA-seq transcriptome profiling

Total RNAs from cultured mouse WT and CD38 KO astrocytes at 5 DIV were used for RNA library preparation using TruSeq Stranded mRNA Sample Preparation Kit (Illumina, Inc., San Diego, CA, USA), with polyA selection for ribosomal RNA depletion.

The RNA-seq libraries were generated in duplicate from 500☐ng of total RNAs extracted from cultured mouse astrocytes. The libraries were sequenced on Illumina HiSeq 2000 to obtain paired-end 101☐bp reads for each sample.

RNA-seq reads from mouse cultured astrocytes were aligned to their reference genomes using the STAR v2.7.0f (56). The Ensembl reference genome used was GRCm38.99 (57). The gene expression profiles were quantified using Cufflinks v2.2.1 (58) from the aligned RNA-Seq reads with gene annotation information corresponding to the above reference genome. Gene expression levels were given by FPKM, which is normalized by the number of RNA fragments mapped to the reference genome and the total length of all exons in the transcript. For bioinformatic analysis of RNA-seq, gene ontology analysis was performed by creating a pre-ranked list of all detected transcripts, ranked by log_2_ fold change < -0.57 and *p* value < 0.05. DAVID, the Database for Annotation, Visualization and Integrated Discovery v6.7 was used to analyze significant changes by KEGG. Significance of enrichment in KEGG pathway analyses was calculated using the p value function given from a modified Fisher’s exact test by the DAVID database.

### Statistical analyses

All data are expressed as mean ± standard error of the mean (s.e.m.), with the number of experiments indicated by (n). Analyses used include unpaired Student’s t-test, one-way ANOVA, or two-way ANOVA and appropriate post hoc analyses (indicated in Figure Legends). Differences were considered statistically significant at a *P* value of <0.05.

## Acknowledgements

We thank Magdalena Götz in GSF-National Research Center for Environment and Health and Mr. Takashi Tamatani and Ms. Nahoko Okitani for providing Cd38^flox/flox^ mice, Glast^CreERT2^ mice and technical assistance, respectively. This work was supported by Grant-in Aid for Scientific Research (21K06407, 18KK0435 for TH, 18KK0255, 18K06500 for OH, 20K09343 for HI and 21K06406, 18K06463 for MT) from the Ministry of Education, Science, Technology, Sports and Culture of Japan, and by Kanazawa University SAKIGAKE Project 2018 and the CHOZEN project.

