## Supplementary Figure for "Astroglial CD38 regulates social memory and synapse formation through SPARCL1 in the medial prefrontal cortex"

**Supplementary Figures and Legends**

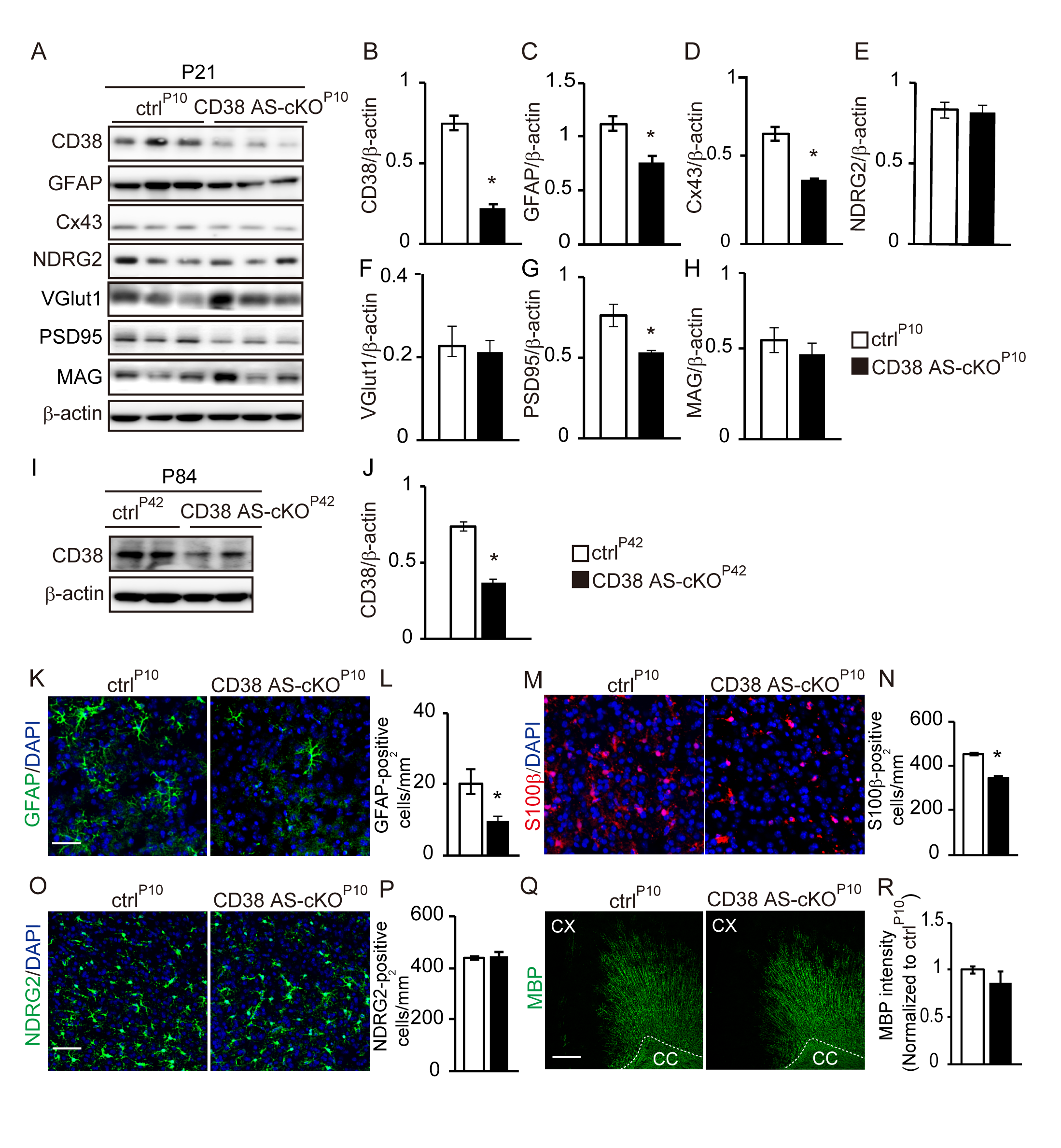

**Supplementary Figure 1. Expression of GFAP, s100β and Cx43 is decreased in the mPFC of CD38 AS-cKO^P10^ mice** (A) Western blot analyses of CD38, GFAP, Cx43, NDRG2, VGlut1, PSD95, MAG and β-actin in the mPFC of ctrl^P10^ and CD38 AS-cKO^P10^ mice at P21. (B-H) Bar graphs depict the relative optical density of proteins normalized to the loading control β-actin. *n* = 5 animals per genotype. Data represent means ± SEM. *P* values were determined by Student’s unpaired *t*-test. ^*^*p* < 0.05 vs ctrl^P10^. (I) Western blot analyses of CD38 and β-actin in the mPFC of ctrl^P42^ and CD38 AS-cKO^P42^ mice at P84. (J) Bar graphs depict the relative optical density of proteins normalized to the loading control β-actin. *n* = 3 animals per genotype. Data represent means ± SEM. *P* values were determined by Student’s unpaired *t*-test. ^*^*p* < 0.05 vs ctrl^P42^. (K, M, O and Q) Immunohistochemistry for GFAP, s100β, NDRG2 in the mPFC and MBP in the motor cortex of ctrl^P10^ and CD38 AS-cKO^P10^ mice at P21. Nuclei were counterstained with DAPI. Scale bars = 50 μm (K, M and O) and 100 μm (Q). (L, N and P) Bar graphs depict the number of GFAP-, s100β and NDRG2-positive cells in the mPFC of ctrl^P10^ and CD38 AS-cKO^P10^ mice at P21. *n* = 4 animals per genotype. Data represent means ± SEM. *P* values were determined by Student’s unpaired *t*-test. ^*^*p* < 0.05 vs ctrl^P10^. (R) Quantification of MBP fluorescence intensity in the motor cortex of ctrl^P10^ and CD38 AS-cKO^P10^ mice at P21. *n* = 4 animals per genotype. Data represent means ± SEM.

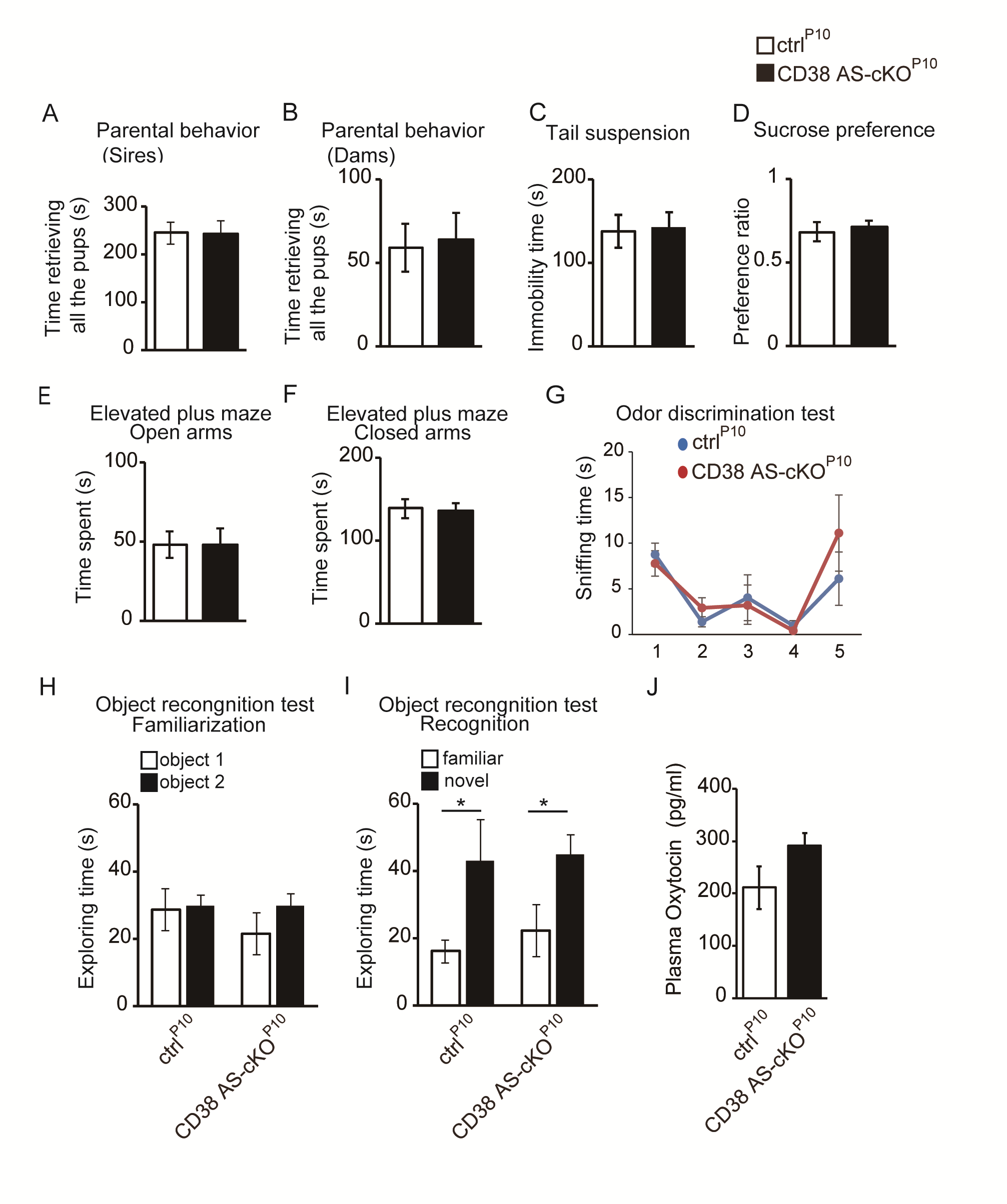

**Supplementary Figure 2.** **Behavioral phenotypes except social memory were not impaired in CD38 AS-cKO^P10^ mice** (A, B) Pup retrieval behavior. Time to complete retrieving all the pups of ctrl^P10^ and CD38 AS-cKO^P10^ sires (A) and dams (B). (C) Tail suspension test. The immobility time of ctrl^P10^ and CD38 AS-cKO^P10^ mice. (D) Sucrose preference test. The preference for 1% sucrose solution consumption by ctrl^P10^ and CD38 AS-cKO^P10^ mice. (E, F) Elevated plus maze test. The time in open arms (E) and closed arms (F) for ctrl^P10^ and CD38 AS-cKO^P10^ mice. (G) Odor discrimination test. Odor habituation (the same odor stimulation repeated four times with 2 min interval, 1st-4th trial) and dishabituation (novel odorant on the 5th trial) in ctrl^P10^ and CD38 AS-cKO^P10^ mice. (H and I) Object recognition test. Exploring time during the acquisition phase (H) and the recognition trial (I). (J) Levels of plasma oxytocin concentration in ctrl^P10^ and CD38 AS-cKO^P10^ mice*. n* = 7 to 10 animals per genotype. Data represent means ± SEM. *P* values were determined by Student’s unpaired *t*-test or two-way ANOVA followed by Scheffe’s *F* test. ^*^*p* < 0.05 between familiar and novel object.

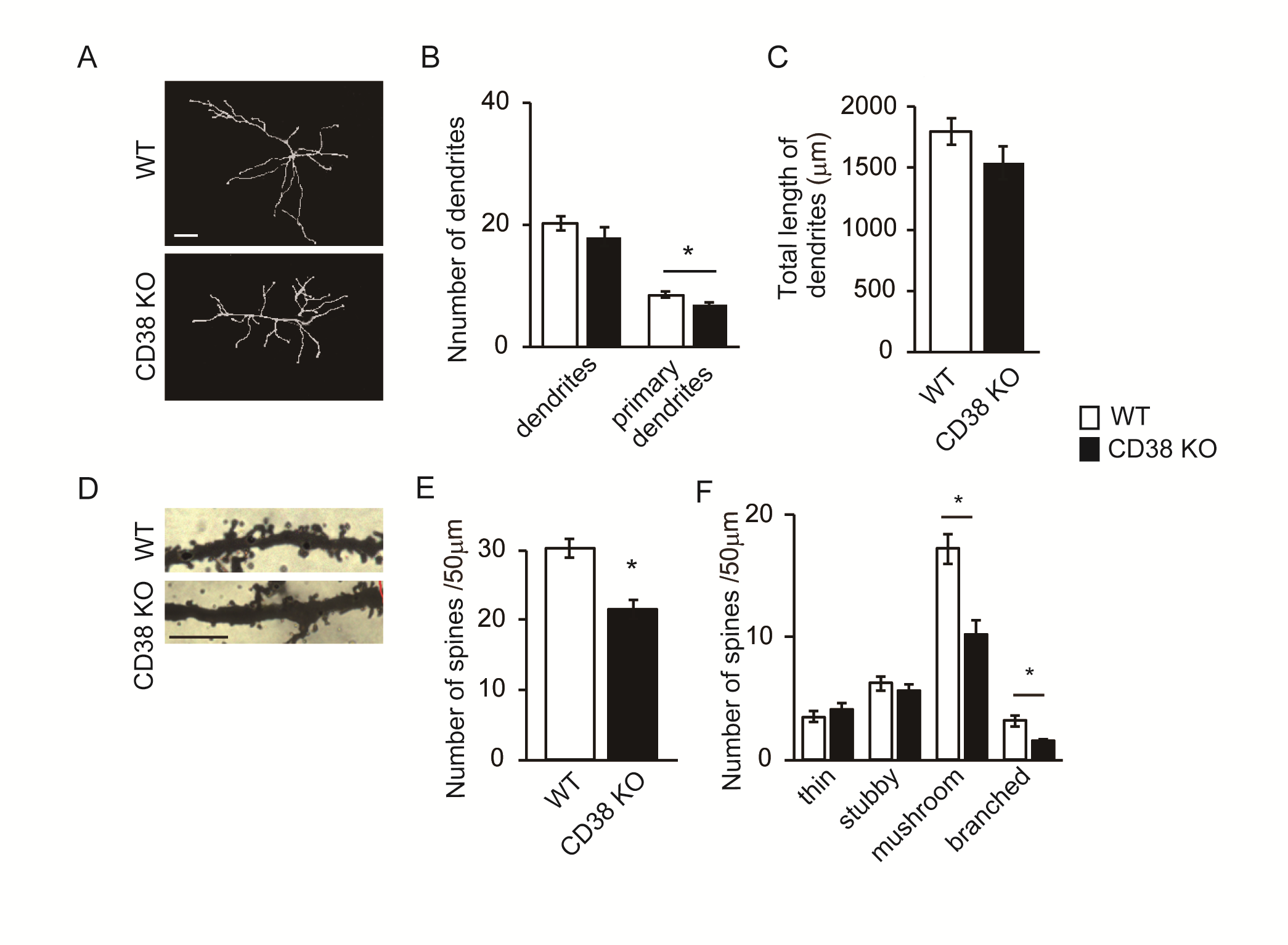

**Supplementary Figure 3. Morphology of dendrites and spines is altered in the mPFC pyramidal neurons in constitutive CD38 KO mice** (A) Images of tracing of Golgi-stained pyramidal neurons in the mPFC of WT and CD38 KO mice. Scale bar = 20 μm. (B) Quantification of the total number of dendrites and number of primary dendrites in WT and CD38 KO mice. (C) Measurement of the total length of dendrites and primary dendrites in WT and CD38 KO mice. (D) Images of Golgi-stained pyramidal neurons in the mPFC of WT and CD38 KO mice. Scale bar = 5 μm. (E) Quantification of the total number of spines on apical dendrites of WT and CD38 KO mice. (F) The percentage of branched, mushroom, stubby and thin-type spines in cortical apical dendritic spines. *n* = 5 cells (B and C) or 4 dendrites (E and F) per animal, 4 animals per genotype. Data represent means ± SEM. *P* values were determined by Student’s unpaired *t*-test. ^*^*p* < 0.05 vs WT.

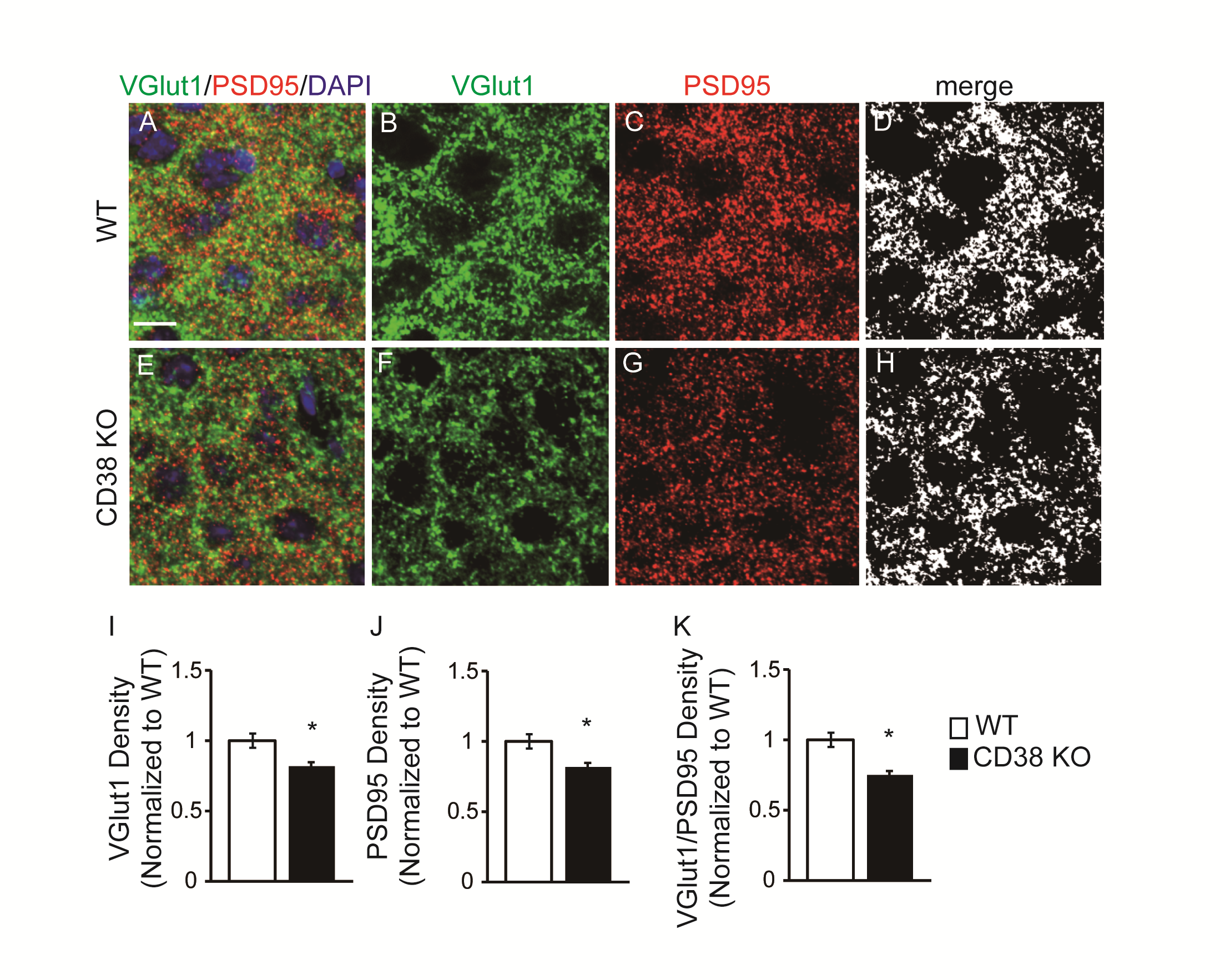

**Supplementary Figure 4. Excitatory synapses are reduced in the mPFC cortical pyramidal neurons in constitutive CD38 KO mice** (A-H) Immunohistochemistry for VGlut1 (green) and PSD95 (red) in the mPFC of WT and CD38 KO mice at P70. Images are single planes from confocal z-stacks. The VGlut1 and PSD95 channels are separated in panels B, C, F and G. Panels D and H are colocalized VGlut1 and PSD95 puncta. Nuclei were counterstained with DAPI. Scale bar = 10 μm. (I-K) Quantification reveals a significant decrease of VGlut1, PSD95 expression and synapses (colocalization of VGlut1 and PSD95) in the mPFC within CD38 KO mice as compared to WT mice. *n* = 4 animals per genotype. Data represent means ± SEM. *P* values were determined by Student’s unpaired *t*-test. ^*^*p* < 0.05 vs WT.

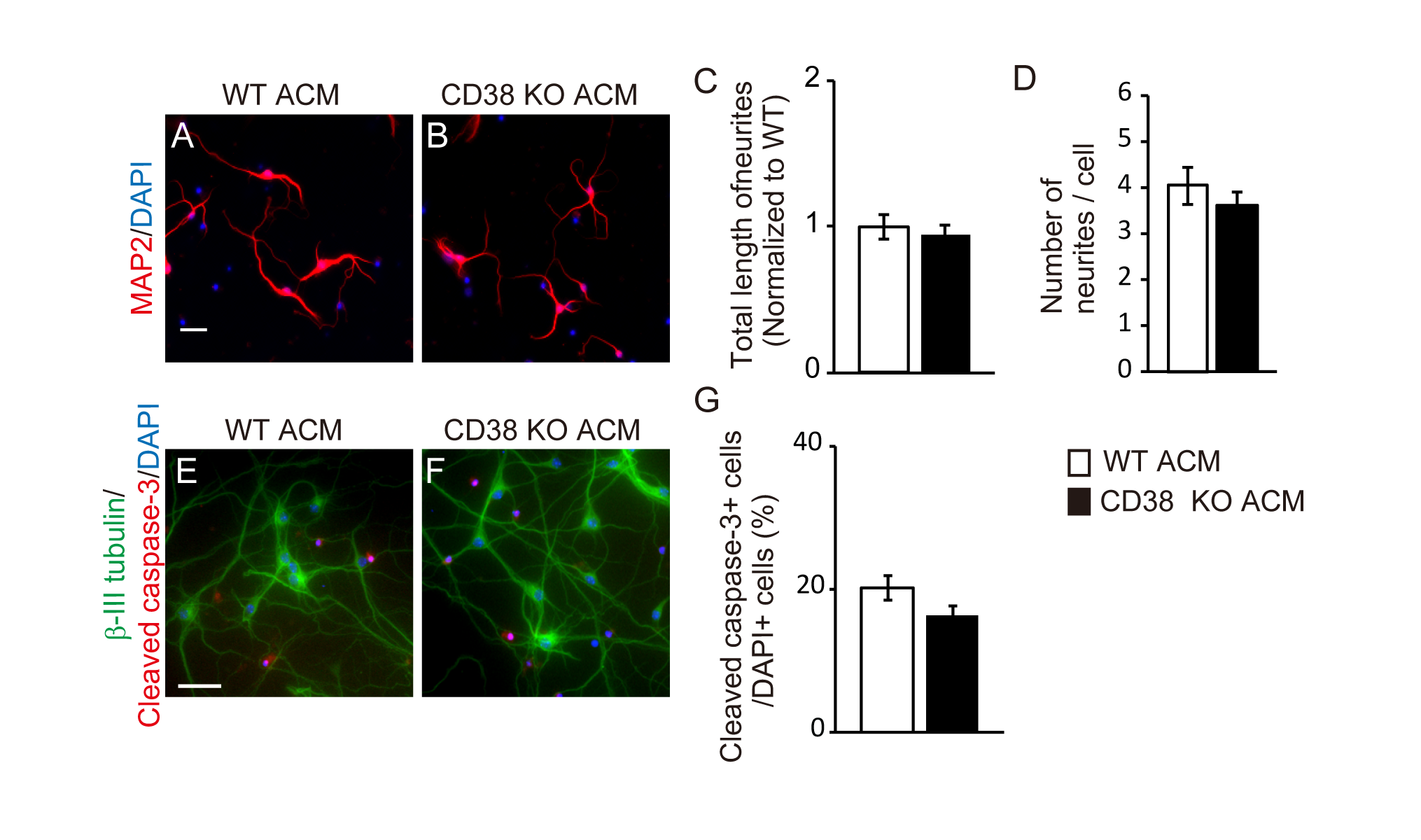

**Supplementary Figure 5. CD38 KOACM does not affect neurite outgrowth and cell viability of cortical neurons** (A, B) Representative images of cortical neurons cultured in ACM of WT or CD38 KO mice at 6 DIV. Cells were treated with WT or CD38 KO ACM from 4 DIV to 6 DIV. Cells were stained with antibody against MAP2 (red). Nuclei were counterstained with DAPI. Scale bar = 50 µm. (C, D) Total length of neurites and number of neurites per cell were measured. *n* = 6 independent cultures from 6 mice per genotype. (E, F) Representative images of cortical neurons cultured in ACM of WT or CD38 KO mice at 6 DIV. Cells are stained with antibody against β-III tubulin (green) and Cleaved caspase-3 (red). Nuclei were counterstained with DAPI. Scale bar = 50 µm. (G) Cleaved caspase-3 positive cells out of DAPI positive cells were counted. *n* = 6 independent cultures from 6 animals per genotype. Data represent means ± SEM.

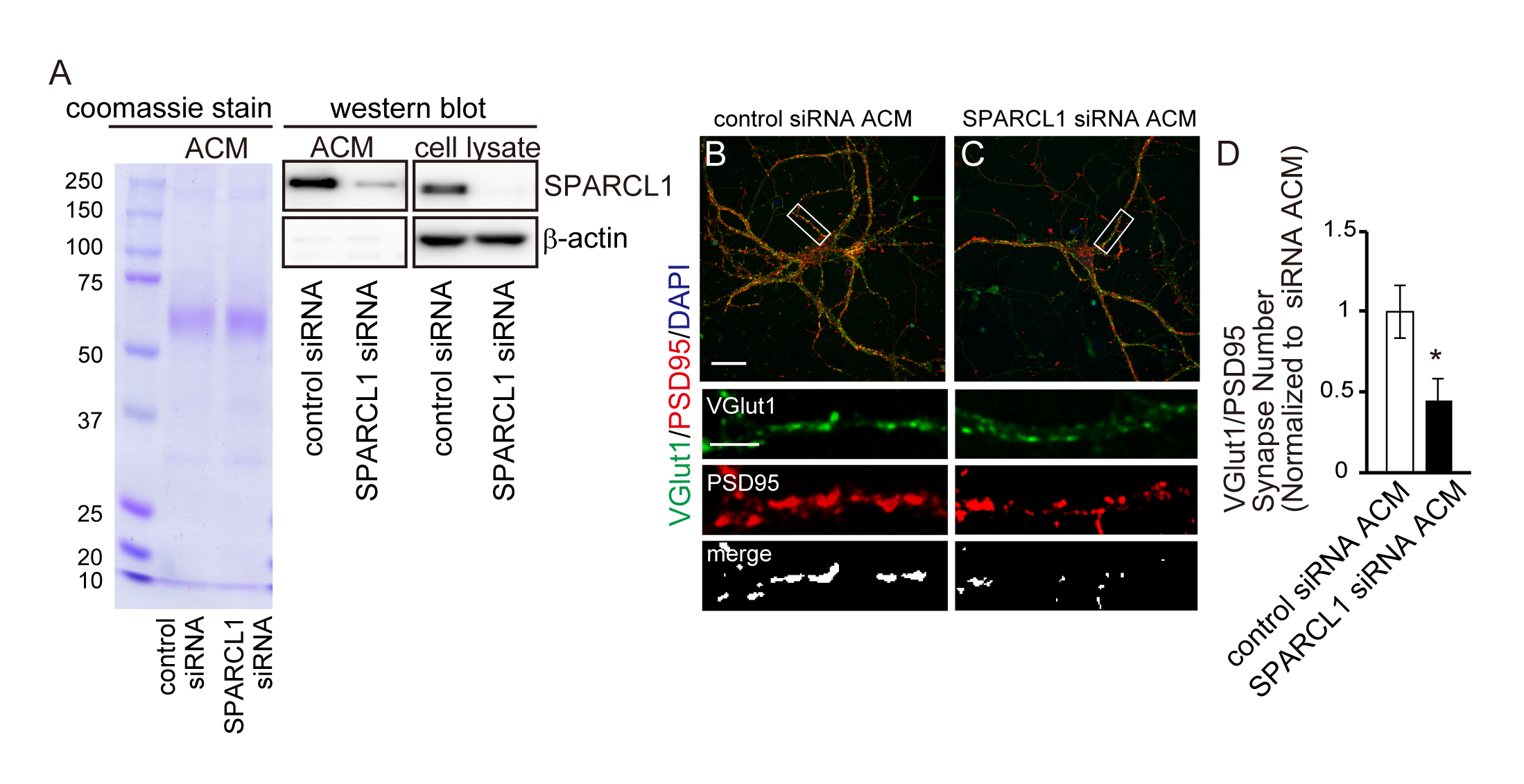

**Supplementary Figure 6. ACM of SPARCL1-knockdown astrocytes reduces synapse formation of cortical neurons** (A) Coomassie stain shows the total protein composition of ACM. Western blot analyses of SPARCL1 and β-actin in ACM and cell lysate of astrocytes transfected with control or SPARCL1 siRNA. (B, C) Representative images of cortical neurons at 14 DIV cultured in ACM of control or SPARCL1 siRNA-transfected astrocytes. Neurons were treated with ACM from 4 DIV to 14 DIV. Insets show individual channels for VGlut1 (green) and PSD95 (red) staining, as well as the merged image. Nuclei were counterstained with DAPI. Scale bar = 20 µm (main image) and 10 µm (inset). (D) Quantification reveals a significant decrease of synapses in cortical neurons cultured in ACM of SPARCL1 siRNA-transfected astrocytes. *n* = 20 cells per condition from 3 independent culture. Data represent means ± SEM. *P* values were determined by Student’s unpaired *t*-test. ^*^*p* < 0.05 vs. control siRNA ACM.

**Supplementary Materials and Methods**

**Parental Retrieval Test**

The design of the experiments for parental retrieval behavior was described previously (1). Virgin males and females of identical genotypes were paired at 56–64 days. A single male and a single female were continuously housed together in a standard mouse maternity cage from the mating period to the delivery of pups and then to postnatal days 3–5. All family units consisted of a new sire and dam, and their first litter of each genotype was used. Thirty minutes before starting the experiment, the cages with the families were placed in the experimental room for habituation. The sire and dam were placed in a new clean cage with new woodchip bedding for 10 min, while the pups were left in the nest in the original cage. Five pups were randomly selected from the litter and placed individually at a site remote from the nest in the original cage. The sires and dams were returned to the original home cage in the presence of their five biological pups to assess parental behavior. Parental retrieval behavior was evaluated by observing the parent behavior for 10 min following the reunion. Time that each sire or dam returned all the pups completely to the nest was measured. The behavioral tests were carried out in a randomly mixed sequence of the experimental groups.

**Tail Suspension Test**

The tail suspension test method was previously described (1). After 30 min of habituation in the experimental room, the mice were hanged by fixing their tails by tape to the suspension bar in a plastic suspension box (55 cm height × 60 cm width × 11.5 cm depth). To prevent observing or interacting with each mouse, the mouse was separated with walls but was not able to contact or touch to the walls. Behavior was recorded for 6 min on video-source and was analyzed using ANY-Maze behavioral tracking software (Stoelting Co.). Total immobility time during the last 4 of the 6 min was used.

**Sucrose Preference**

Experimentally naïve, young adult, male and female mice (6–8 weeks old) were housed individually and were given a two-bottle choice between distilled water and sucrose solution at a 1% concentration based on our previous results (1), which were both available ad libitum. The bottle positions remained constant. Fresh sucrose solution was prepared each day. Cumulative water and sucrose intakes during 24 h were calculated by weighing. Food was provided ad libitum, but food intake was not recorded in this experiment. Sucrose preference was calculated according to formula W0 − W1/((W0 − W1) + (S0 − S1). Where S0 and W0 are the weight of bottles with sucrose solution

and water before the test and S1 and W1 after 24 h after the test.

**Elevated plus Maze**

This test was based on a method that was described previously (2). The maze was made of four black plexiglass arms with two open arms (67 × 7 cm) and two walled arms (67 × 7 × 17 cm) facing each other and connected by a neutral space in the center at 55 cm above the floor under dimly illuminated light (20 lux). The camera-assisted ANY-Maze software automatically tracked the time spent, frequency of entry into each arm in 5 min. Time spent in open, and close arms was evaluated.

**Object recognition test**

The novel object recognition test was conducted in a big soundproof box (60 cm × 60 cm × 100 cm), containing a square, high-walled arena (25 cm × 25 cm × 25 cm) at its bottom. Briefly, it consisted of three phases: habituation sessions (day 1 and day 2, no object inside, 20 min/day), training session (day 3, two identical objects placed symmetrically, 10 min), and test session (one familiar object replaced by a novel one, 5 min, session was started immediately after replacing the objects). The two kind of objects used in the test have similar size and smell but differ in shape and texture. The camera-assisted ANY-Maze software automatically tracked the time spent near the objects. The time spent exploring each object was scored when the subject was sniffing towards the object within 2 cm.

**Odor discrimination test**

The olfactory habituation/dishabituation test was performed as described previously. All the test mice were single housed before the test. During the test mouse was put in Plexiglas box (90 cm × 50 cm × 30 cm). Test session consisted from 5 trials (each for 5 minutes) with 5 minutes paused between them, when mouse was returned to home cage. During sessions 1-4 cotton pad containing neutral odorant (124-03892, R(+)-Limonene, WAKO) was presented. The test mice were allowed to investigate odor. In session 5 another neutral odorant (150-00134, 1-Octanol, WAKO) was presented. Time spent sniffing the odor was measured by manual observation with a stopwatch. Time was only scored when the test mouse was sniffing the swab within 2 cm.

**Oxytocin measurement**

Thoracotomy was performed under anesthesia by isoflurane, and blood sample was obtained from the ventricle by cardiac puncture. An Oxytocin ELISA kit (ADI-901-153, Enzo Life Sciences, New York, NY, USA) was used for oxytocin analysis by following manufacture’s protocol.

**siRNA transfection**

We obtained small interference RNA (siRNA) from Sigma. The sequence of the mouse SPARCL1 targeted was as follows: 5′-CTATTCCTGCTTGTACGG-3′　(SASI_Mm01_00135946). We used Stealth RNAi™ siRNA negative control med GC (Thermo Fischer SCIENTIFIC) as a control siRNA. Transfection of siRNA was performed using RNAimax (Invitrogen) according to the manufacturer's instructions.
